# Vibrotactile transcutaneous auricular vagus nerve stimulation (taVNS) during working memory: convergent electrophysiological evidence for cortical efficiency

**DOI:** 10.64898/2026.10.07.756069

**Authors:** Gansheng Tan, Phillip Demarest, Tatiana Kardashina, Josh Adams, Jon T. Willie, Jarod Roland, Jenna L. Gorlewicz, Peter Brunner, Eric C. Leuthardt

## Abstract

**Background:** Working memory, the cognitive ability to maintain and manipulate a finite amount of information, is fundamental in reasoning, learning, and decision-making. Working memory erodes in conditions from Alzheimer’s disease to stroke to healthy aging. Transcutaneous auricular vagus nerve stimulation (taVNS) is a promising neuromodulation method for improving working memory. By stimulating the auricular branch of the vagus nerve, taVNS is thought to engage ascending arousal systems such as the locus coeruleus-norepinephrine (LC-NE) system. However, the physiological link between LC-NE system modulation via taVNS and improved working memory remains largely unknown. This mechanistic gap limits the rational optimization of the stimulation protocol and the identification of the cognitive functions and patient populations that are likely to benefit.

**Objective:** We test two competing hypotheses for how taVNS may improve working memory: sensory amplification or enhancement of cortical efficiency. The sensory-amplification hypothesis predicts increased cortical responses to stimuli, whereas the cortical-efficiency hypothesis predicts reduced cortical responses.

**Methods:** Twenty healthy adults completed N-back working memory tasks during three counterbalanced sessions: baseline, active vibrotactile taVNS (vtaVNS), and sham stimulation, while we simultaneously recorded electroencephalography, electrocardiography, eye-tracking, and behavioral responses. To study the effect of vtaVNS on neuronal activity, we also recruited 20 patients with epilepsy undergoing invasive monitoring. Of these, 19 fully completed the spatial working-memory task. We quantified cortical responses to visual stimuli using event-related potentials (ERPs) in scalp EEG and high-gamma activity in intracranial local field potentials. To examine prediction of the cortical-efficiency hypothesis, we further explored the effect of vtaVNS on distance to criticality and heartbeat-evoked potentials.

**Results:** Active vtaVNS reduced both scalp-recorded visual ERP and high-gamma response in temporal cortices, relative to sham stimulation. Moreover, smaller visual event-related potentials were associated with successful encoding during the most demanding working-memory condition (i.e., 4-back task). Exploratory analyses showed that vtaVNS shifted cortical dynamics closer to criticality.

**Conclusion:** Our results converge on the conclusion that vtaVNS does not uniformly amplify cortical responses to external stimuli. Instead, vtaVNS enhances cortical efficiency, supporting more effective allocation of cognitive resources between exteroceptive and mnemonic processes. These findings provide a framework for predicting when taVNS will be most beneficial and for optimizing stimulation protocols across cognitive and clinical applications.

**Highlights:**

- Consistent visual event-related potential in human subjects during an N-back working memory task
- Vibrotactile taVNS reduced stimulus-evoked cortical responses in both scalp EEG and intracranial high-gamma activity (70-150 Hz).
- Reduced visual event-related potentials are associated with successful encoding in the 4-back task.
- Vibrotactile taVNS shifted cortical dynamics toward a near-critical regime.

## Introduction

Working memory, the ability to maintain and manipulate sensory information over short intervals, is key to intelligent behaviors, including reasoning, planning, and decision-making. Yet working memory is fragile. It declines with age and is disrupted by a broad range of neurological conditions, including Alzheimer’s disease, stroke, and epilepsy^1–3^. Existing approaches to restoring or protecting working memory remain limited. Pharmacological agents targeting the dopamine or noradrenergic systems act diffusely and continuously across the body, with modest or inconsistent cognitive effects^4,5^; and cognitive training programs such as memory games rarely produce significant improvement^6^. Thus, there is a need for more effective interventions to improve working memory.

Transcutaneous auricular vagus nerve stimulation (taVNS) offers a novel approach to improving memory because (1) it innervates parts of the external ear, allowing for easy access and non-invasive activation; and (2) the auricular branch of the vagus nerve projects, via brainstem nuclei, to cortical and subcortical regions implicated in memory, including locus coeruleus and hippocampus^7–10^. One form of taVNS delivers electrical pulses to auricular regions innervated by the auricular branch of the vagus nerve, most commonly the cymba conchae^11,12^. More recently, vibrotactile taVNS (vtaVNS) has emerged as an alternative, motivated in part by comparative evidence that the auricular branch of the vagus nerve is evolutionarily related to a mechanosensory system for detecting vibration and water movement in aquatic ancestors^7,9,13,14^. The human auricular branch of the vagus nerve contains myelinated *Aβ* afferent fibers, which primarily transmit non-painful, discriminative light touch and vibration from cutaneous mechanoreceptors^15^. Both electrical taVNS and vtaVNS have demonstrated success in improving memory^14,16–18^. However, the physiological mechanism linking taVNS to memory improvement remains poorly understood, leaving open the question of which patient populations may be most responsive and under what circumstances taVNS achieves the greatest benefit. Understanding how taVNS alters cortical physiology during memory is therefore a prerequisite for its rational development as a memory-improvement intervention.

taVNS engages multiple physiological systems, altering neurotransmitter release, autonomic tone, and inflammatory signaling^19–22^. Among these systems, its cognitive benefits, such as improving working memory, are more closely related to modulation of the locus coeruleus-norepinephrine (LC-NE) system^14,23^. Specifically, taVNS activates the nucleus tractus solitarius (NTS), modulates activity in the LC, which in turn regulates norepinephrine release in cortical and subcortical structures (**Figure 1A)**^11,24^. Most mechanistic evidence to date has been derived from electrical taVNS, and studies using vtaVNS also suggested modulation of the LC-NE pathway and inflammatory signaling^9,14,25^. However, how engagement of the LC-NE pathway reshapes the cortical dynamics that support working memory remains unknown.

**Figure 1.**
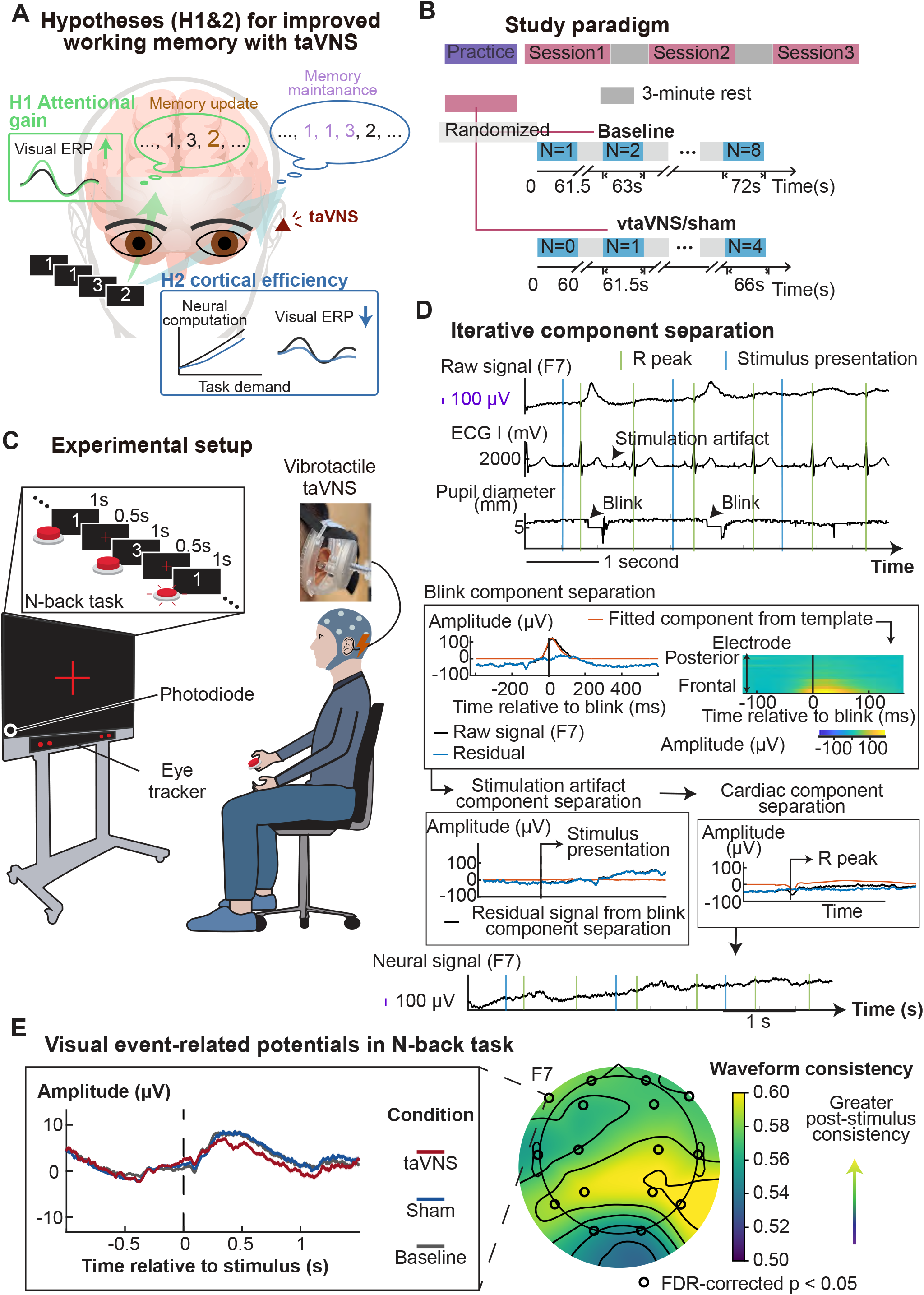
Mechanistic hypotheses and experimental design for studying how taVNS modulates working memory. **(A)** Two competing hypotheses for how taVNS improves working memory. Based on the modulation of the LC–NE system via taVNS, we consider two candidate mechanisms. Under the sensory-amplification hypothesis, taVNS amplifies sensory responses to stimuli, producing larger visual event-related potentials (ERPs) or population activity indices with high-gamma power in local field potentials. This stronger sensory drive would support higher-fidelity encoding of stimuli into working memory and prolong its maintenance. Under the cortical efficiency hypothesis, taVNS reduces the neural computation devoted to processing external stimuli, producing smaller visual ERPs or population activity. Efficient use of cognitive resources would support memory maintenance, updating, and retrieval, thereby improving performance. **(B)** Study paradigm. Each healthy participant completed three sessions of the N-back task, with orders randomized across baseline (device worn, no stimulation), sham stimulation, and active vtaVNS. To titrate task performance, Subjects completed N-back tasks with N ranging from 1 to 8 in the baseline session. In active or sham stimulation sessions, subjects completed N-back tasks with N ranging from 0 (i.e., attention task) to 4. (**C**) Physiological recording setup during the N-back task. Participants viewed a sequential stream of digits (drawn from 1, 2, and 3) and pressed a button whenever the current digit matched the one presented N trials earlier. We recorded scalp EEG, electrocardiogram, gaze position, pupillometry, and behavioral responses simultaneously. (**D**) Iterative component separation from EEG signals. We used a template-matching approach to sequentially remove oculomotor (blink), stimulation-artifact, and cardiac components from the EEG. We first constructed a template for the blink component, fit it to the raw signal, and subtracted it to obtain a blink-corrected residual. We then constructed a template for the stimulation artifact, fit it to the residual, and subtracted it; the same procedure was repeated for the cardiac component. The neural component (bottom) is the final residual signal after all three artifactual components have been sequentially identified and removed. (**E**) Visual ERP during the N-back task. The topographic map shows the trial-to-trial consistency of EEG signal following visual stimulus across the scalp. The adjacent trace shows the visual ERP time course for an exemplar electrode.

The LC-NE system regulates stimulus responsiveness and prioritization of task-relevant information^26–29^. LC neurons operate through interacting tonic and phasic modes: tonic activity sets the background level of cortical arousal, whereas phasic bursts transiently release norepinephrine in relation to behaviorally relevant events^23^. Through these actions, LC–NE signaling can alter both the responsiveness of cortical circuits to sensory input and the cortical state within which limited processing resources are allocated^26,27,29^. We considered two mechanistic accounts of how taVNS may improve working memory: a sensory amplification account and a cortical-efficiency account. Under the sensory-amplification account, taVNS increases neuronal sensitivity, thereby increasing the fidelity and persistence of the representation of incoming sensory information in memory^27,30,31^. This account predicts larger cortical responses to stimuli and the greatest cognitive benefit of taVNS when performance is limited by insufficient sensory evidence, such as when detecting subtle visual changes. Under the cortical-efficiency account, taVNS promotes an efficient state in which neural computation is flexibly prioritized according to behavioral goals^32,33^. The cortical-efficiency account predicts smaller cortical responses with preserved performance, and the greatest benefit when task demands strain limited cognitive capacity, such as under high working-memory load or when distracting sensory input competes with mnemonic processing^34^. Adjudicating between these accounts therefore informs therapeutic implications for taVNS.

We previously demonstrated that 6 Hz vtaVNS improves working memory performance selectively under high cognitive load, which led us to hypothesize that vtaVNS modulates cortical efficiency^35^. We initially tested whether vtaVNS is associated with a reduction in cortical response to sensory input. Although such a reduction is consistent with the cortical-efficiency hypothesis, it is not sufficient evidence on its own, because reduced cortical responses could reflect reduced attention^36^. We therefore tested two further predictions that, together with reduction in cortical response, would distinguish cortical efficiency from disengagement. We assessed, first, whether the reduction in cortical response is associated with successful memory encoding, particularly under high working-memory load; and second, whether vtaVNS shifts cortical dynamics toward a critical operating regime, in which computation can be flexibly allocated according to behavioral goals^37^. Criticality describes the boundary between states in which activity is rapidly damped (subcritical dynamics) and states in which activity propagates excessively through the network (supercritical dynamics). Cortical networks operating near this boundary are thought to support the flexible transmission and reconfiguration of information in response to changing demand^38^.

To evaluate these predictions, we measured cortical responses at complementary spatial and physiological scales. At the macroscopic scale, we quantified visual event-related potentials (ERPs) with scalp electroencephalography (EEG), isolating neural signals from ocular, cardiac, and stimulation-related artifacts through simultaneous eye-tracking and electrocardiography with a component separation approach^39,40^. Scalp EEG mixes contributions from multiple cortical sources. To strengthen both the anatomical and physiological interpretations of the effects of vtaVNS on visual ERP, we applied source localization to identify cortical regions that could account for the vtaVNS-related change in the visual ERP. To test these anatomical hypotheses, we examined high-gamma activity (70–150 Hz) in stimulus-related intracranial local field potentials (LFPs), which index local population-level neuronal firing^41,42^. Across scalp EEG recordings from 20 healthy participants and intracranial EEG recordings from 20 patients with epilepsy, we found reductions in both visual ERPs and stimulus-related high-gamma activity in the active vtaVNS condition compared to the sham control. Separately, smaller visual ERP responses were associated with successful memory encoding under high working-memory load. vtaVNS also reduced the distance to criticality, indicating that cortical dynamics shifted toward a more critical operating regime^38^. This converging evidence supports a model in which vtaVNS improves working memory not by amplifying sensory responses, but by shifting the brain toward a critical state that allocates limited processing capacity more efficiently.

## Results

Twenty healthy adults completed N-back tasks at varying levels of difficulty across three randomized-order sessions (active vtaVNS, sham, and baseline, **Figure 1B**). We also recruited 20 neurosurgical patients undergoing invasive monitoring for seizure-onset-zone localization, of whom 19 completed the entire spatial working-memory task. The study was approved by the Washington University in St. Louis Institutional Review Board (IRB ID#: 202101161). Written informed consent was obtained from all study participants. The healthy population received 6 Hz vtaVNS delivered to the left concha or earlobe during active vtaVNS and sham sessions, respectively. Vibrotactile stimulation was paired with stimulus presentation and delivered using a customized device that houses a 5 mm eccentric rotating mass motor and is secured to the ear with an adjustable contact depth and an elastic head strap^14^ (**Figure 1C**). Analysis of behavioral data confirmed that healthy subjects performed the n-back task as instructed^14^. Neurosurgical patients also successfully performed the spatial working-memory task, with a mean accuracy of 78.7% (SD 16.0%), supporting that they remained engaged with the task and encoded the presented visual information (see Supplementary Material for exploratory analyses of behavioral performance across active vtaVNS and sham sessions). The present study focused on the electrophysiological mechanism of vtaVNS during the working memory task.

### Robust visual event-related potentials during N-back tasks

Visual ERP analysis requires separating neural activity from physiological and stimulation-related artifacts. Standard artifact-removal methods, such as independent component analysis, assume that artifactual and neural signals are statistically independent^43^. When this assumption is violated, independent component analysis may incompletely separate artifacts or distort neural signals. In our study, this assumption is difficult to satisfy because visual ERPs may be temporally related to blinking following the presentation of a visual stimulus. We therefore decomposed our EEG recordings into four parsimonious components: blink component, stimulation artifact component, cardiac component, and neural component (**Figure 1D**). We estimated artifact components from event-locked templates constructed around blinks, stimulation onsets, and R-peaks (i.e., the upward deflection of the QRS complex on an electrocardiogram), and then fit them to the EEG data, allowing bounded variation in amplitude and timing (see **Methods: Supervised sequential artifact-component separation**). We separated components sequentially, with the blink component first, followed by stimulation and cardiac artifacts. We used the remaining signal as the neural component for subsequent visual event-related analysis. We verified that our template-matching algorithm separated components as expected using both quantitative measures and qualitative visual inspection of waveform shape. If the fitted template includes a substantial artifact component, subtracting it should reduce the signal energy. Little or no change in energy may occur when the artifact is weak relative to neural activity, whereas a substantial increase in energy suggests that the fitted component introduces distortion. Across subjects and sessions, artifact component subtraction reduced signal energy within the artifact window (**Fig. S1B**, median change in signal energy (Δ*E*) = 0.411, 95% confidence interval (CI) = [0.354, 0.462], N = 57, for blink component; median Δ*E*= 0.055, 95% CI = [0.020, 0.097], N = 40, for stimulation artifact component; median Δ*E* = 0.092, 95% CI = [0.020, 0.112], N = 60, for cardiac component). The fitted template parameters, including scaling and temporal shift, indicate stable artifact morphology. (**Fig. S1C–D, Supplementary Materials**). Together, the template-matching procedure separated the neural signal from the blink, stimulation, and cardiac artifact components, providing the basis for the subsequent analyses of visual ERPs.

We next confirmed that our electrophysiological data reproduced the previous observation of visual ERP in EEG^44,45^. In the presence of consistent visual ERPs, we expect similar waveforms following stimulus presentation^45^. To test this, we compared waveform consistency before and after stimulus presentation. We found that the visual ERPs across all recording electrodes were significantly more consistent than the field potential before stimulus presentation (**Figure 1E, Fig. S1D**, median area under the ROC curve based on inter-trial correlation = 0.575; all False Discovery Rate (FDR)-corrected p < 0.001; linear mixed-effects model).

Our data demonstrate robust visual event-related potentials across the brain during the N-back task.

### vtaVNS reduces visual event-related potentials

To evaluate the first prediction of the cortical-efficiency hypothesis, we test whether vtaVNS is associated with reduced visual ERPs. An increase in visual ERP would instead support the sensory-amplification hypothesis^40^. We compared visual ERPs between active vtaVNS and sham using a cluster-based permutation test. This analysis identifies spatially and temporally consistent changes in visual event-related potential amplitude between active and sham stimulation using a threshold-free cluster enhancement (TFCE) method^46^. We found that the visual ERPs were reduced in the active vtaVNS condition, compared to sham stimulation (**Figure 2A**). The most prominent cluster was observed over temporal-central electrodes from 0.250 to 0.600 s after stimulus presentation (**Figure 2A**, minimum corrected p = 0.008, peak effect = -0.518 z-score unit, *N*_*subject*_ = 20, permutation test).

**Figure 2.**
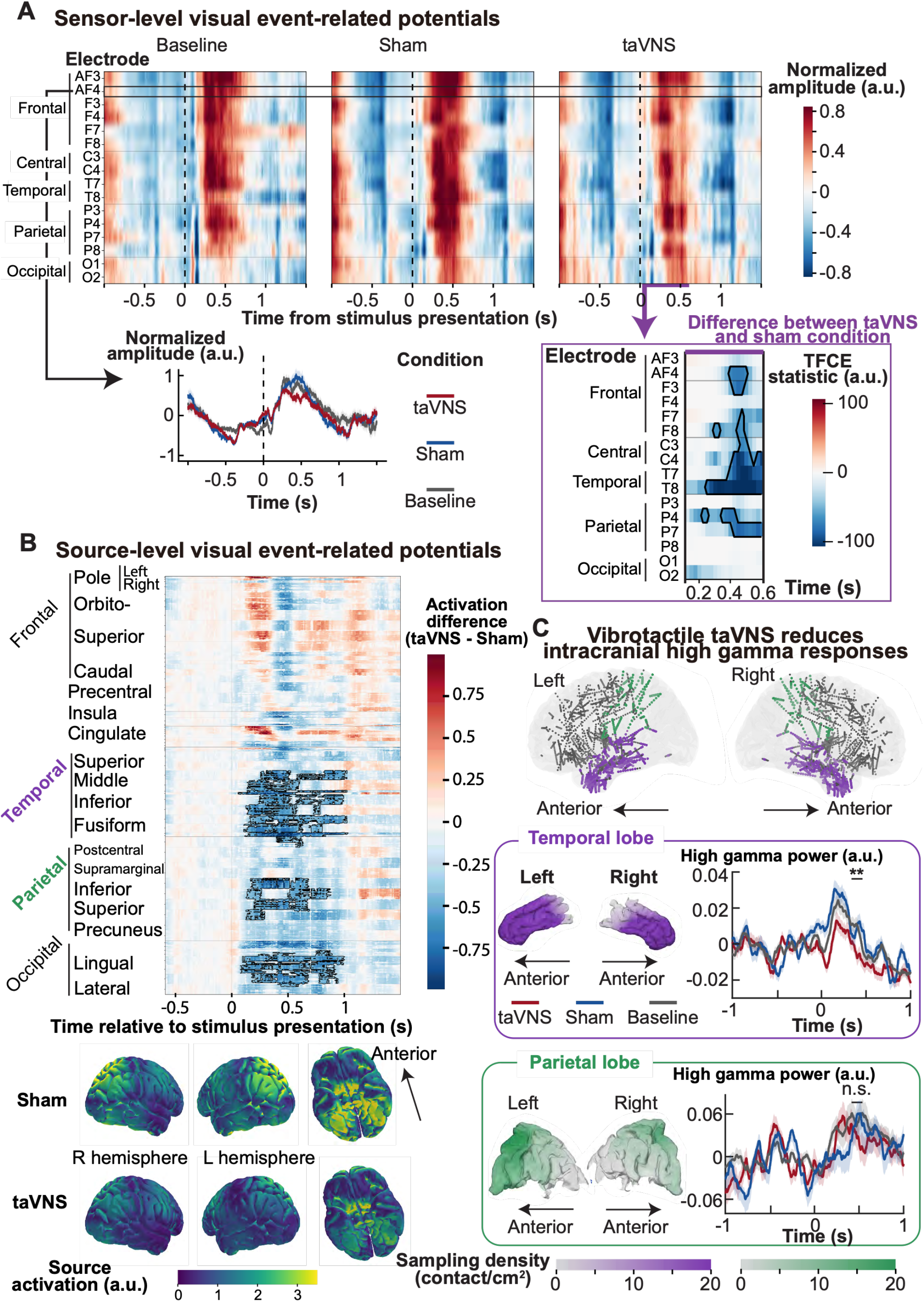
vtaVNS reduces cortical response to visual stimulus. (**A**) Sensor-level visual ERP across baseline, sham, and vtaVNS sessions. Heatmaps show the visual ERP at each electrode (ordered anterior to posterior) over time, with color representing amplitude z-scored to the pre-stimulus baseline period (−0.5 to 0 s). The visual ERP time course across the three sessions is plotted in the bottom left for a representative right-prefrontal electrode. The bottom-right plot with a purple outline depicts the difference in visual ERP amplitude between vtaVNS and sham sessions. Black outlines mark electrode-time clusters showing a significant difference between the vtaVNS and sham conditions, identified using a cluster-based permutation test with threshold-free cluster enhancement. (**B**) Source-level difference in electrical field magnitude between vtaVNS and sham. We estimated cortical source activity using standardized low-resolution brain electromagnetic tomography (sLORETA). The heatmap shows the vtaVNS-minus-sham difference in electrical field magnitude across cortical regions (ordered anterior to posterior) over time; red indicates greater magnitude in the vtaVNS condition, blue indicates greater magnitude in the sham condition. The bottom plot shows the magnitude of the electrical field across the cortical surface for vtaVNS and sham conditions, averaged over 0.2–0.6 s. (**C**) vtaVNS reduces high-gamma activity recorded intracranially. The top plot shows electrode contacts in the temporal lobe (purple) and parietal lobe (green) in a separate cohort of 20 patients with epilepsy undergoing intracranial monitoring. The bottom two plots show high-gamma activity over time for the temporal (purple outline) and parietal (green outline) lobes, across vtaVNS, sham, and baseline conditions. The color intensity of cortical surfaces reflects the density of sampling electrode contacts. Time 0 marks stimulus presentation during the encoding phase of a spatial working memory task.

In an N-back task, each visual item is not merely an external stimulus. It must also be encoded, compared with the item held in memory, and used to update the working-memory state. We next investigate the psychological relevance of the reduction in visual ERP: whether it reflects working memory updating in addition to visual processing. If visual ERP represents a memory update, the relationship between visual ERP amplitude and working memory load should be positive. Our cluster-based permutation test showed no significant effect of task difficulty on visual event-related potentials. Posterior power analysis showed that our data had 86% power to detect a medium correlation (Pearson’s r = 0.3; **Fig. S1E**), suggesting that visual ERP is more closely related to sensory processing than to working-memory updating.

To identify the functional-anatomical brain region that generated the sensor-level reduction in visual event-related potential, we source-localized the visual evoked potential using standardized low-resolution brain electromagnetic tomography (sLORETA)^47^. Following the same reasoning that a true effect of vtaVNS should be temporally and spatially continuous, we applied spatial-temporal coherence weighting on source-level current-field magnitude (See **Methods: Source localization**). The three regions showing most consistent changes in current-field magnitude between vtaVNS and sham are bilateral middle, inferior, and medial temporal lobe (**Figure 2B**, 0.23–0.82 s; mean magnitude difference (a.u.) = -0.722), bilateral superior and inferior parietal lobules (0.23–0.90 s; magnitude difference (a.u.) = -0.680), and bilateral lingual and lateral occipital cortex (0.18–0.64 s; magnitude difference (a.u.) = -0.721, N = 20). These findings affirm the first prediction of the cortical-efficiency hypothesis that vtaVNS reduces cortical activity required for sensory processing.

### vtaVNS reduces intracranial high-gamma responses

Our healthy-subject study generated the hypothesis that vtaVNS reduces visual ERPs in the temporal, parietal, and occipital regions compared to sham. To test whether this reduced current-field magnitude results from reduced population-level neuronal activity, we recorded intracranial local field potentials from 20 neurosurgical patients undergoing invasive monitoring for seizure-onset-zone localization (see **Supplementary Table S1** for patient characteristics). Because the N-back task used in the healthy cohort was not well tolerated by neurosurgical patients, the patient cohort performed a classic spatial working memory task designed to assess the effect of vtaVNS on cortical responses to visual stimuli (see **Methods, Movie S1**). Consistent with our hypothesis, compared to sham stimulation, vtaVNS reduced high-gamma activity, a physiological marker of large-scale neuronal activity, in bilateral middle, inferior, and medial temporal lobe (**Figure 2C**, linear mixed-effect model, *N*_*electrode*_ = 733, *β*_*VNS*_ = -0.018, standard error = 0.006, p = 0.003). vtaVNS also trended toward reducing intracranial high-gamma responses in the superior and inferior parietal lobes; however, this effect was not significant (**Figure 2C**, linear mixed-effect model, *N*_*electrode*_ = 72, *β*_*taVNS*_ = -0.012, standard error = 0.029, p = 0.692). In our cohort of focal epilepsy patients, invasive monitoring did not commonly involve depth electrodes in the occipital cortex, which precluded testing whether vtaVNS also reduced occipital neuronal activity.

### Reduced visual ERP was associated with successful encoding in 4-back tasks

Our electrophysiological data showed that vtaVNS reduced population neural activity following stimulus presentation, suggesting that vtaVNS reallocated cognitive resources from exteroceptive attention to task-relevant neural computation^48^. We next examined the second prediction of the cortical-efficiency hypothesis that the reduction in cortical response to visual stimuli should be associated with successful memory encoding. Thus, we compared visual ERP amplitudes between successful and unsuccessful encoded trials in the 0.5-0.6 s window where the vtaVNS-induced reduction was strongest (**Figure 2A**). In the 4-back task, the visual ERP was significantly smaller for successfully than for unsuccessfully encoded trials (**Figure 3A;** Cohen’s d = -0.520, p = 0.032, permutation test, N = 20). We found no such difference in the easier N-back tasks (**Figure 3B**; Cohen’s d = -0.056, 0.368, -0.300; p = 0.856, 0.116, 0.195 for 1-, 2-, and 3-back tasks, respectively). This finding aligns with previously published behavioral findings that vibrotactile taVNS improved working memory performance on difficult tasks^14^.

**Figure 3.**
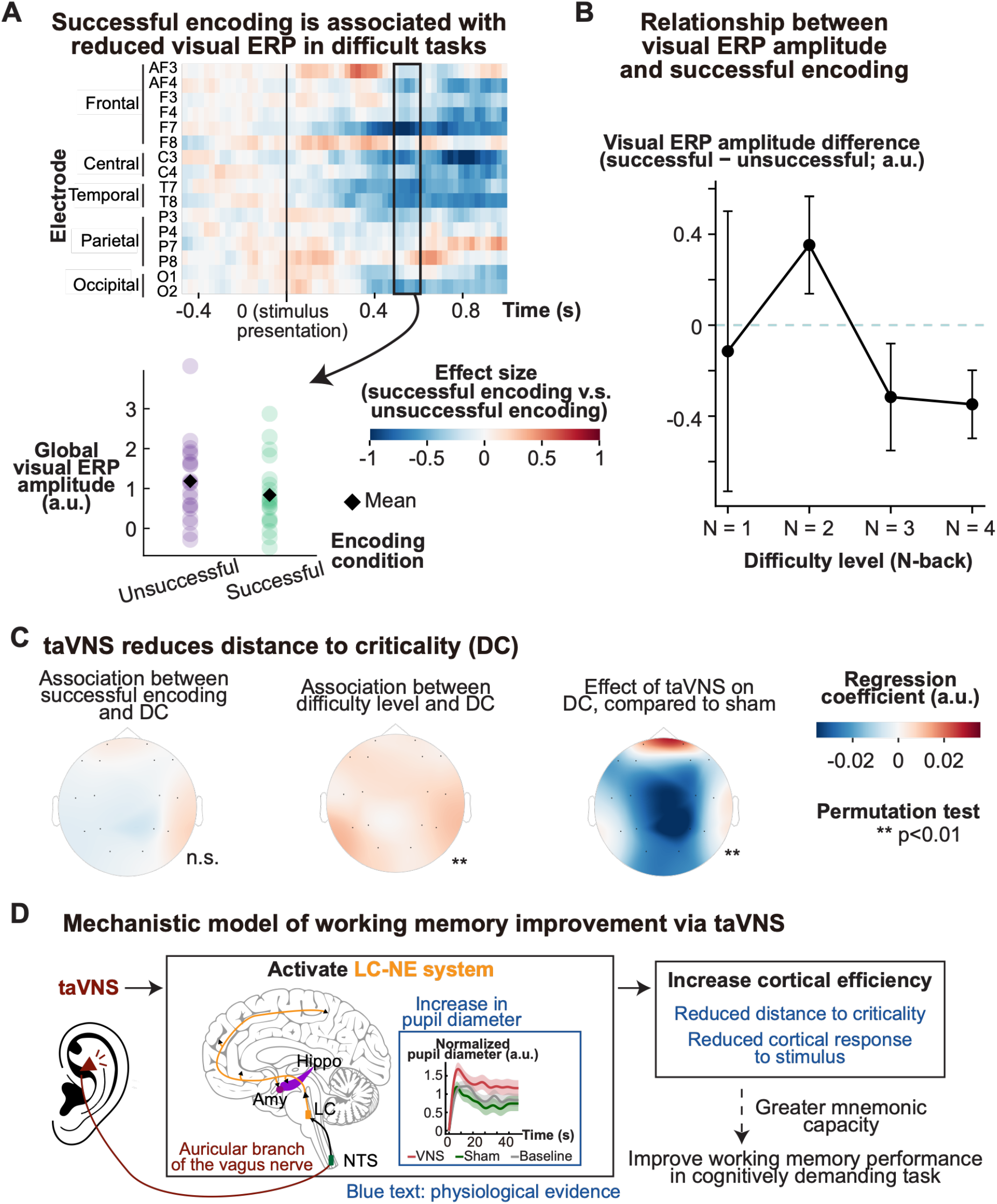
Reduced visual ERPs are associated with successful encoding. (**A**) Difference in visual ERP amplitude between successful encoding and unsuccessful encoding trials in the 4-back task. The top plot is an electrode-by-time heatmap showing successful-minus-unsuccessful visual ERP difference. The bottom plot shows visual ERP amplitude between 0.5–0.6 s following stimulus presentation, averaged across electrodes for successful versus unsuccessful encoding trials. Each participant’s paired values are shown individually. (**B**) Relationship between visual ERP amplitude and successful encoding across working memory task difficulty. The plot shows the mean difference in visual ERP amplitude between successful and unsuccessful encoding at each working-memory difficulty level; error bars indicate ± SEM across participants. (**C**) vtaVNS reduces distance to criticality. The topological graphs visualize the regression of distance to criticality (DC) in a linear mixed-effects model against successful encoding, difficulty level, and stimulation condition. Blue color denotes a negative association. Distance to criticality during baseline, sham, and active vtaVNS sessions is shown in **Fig. S4C**. (**D**) Proposed mechanistic model of working memory improvement via vtaVNS. This schematic synthesizes the behavioral and physiological findings: vtaVNS activates the LC–NE system; this neuromodulation shifts cortical dynamics toward a near-critical operating regime, within which cognitive resources are flexibly and efficiently allocated among perception, memory updating and maintenance, and executive control, thereby improving working memory performance. The pupil-diameter response shown in the schematic is derived from experimental data from our previous publication and is reproduced here to illustrate evidence of LC–NE-related arousal^14^. The dashed arrow indicates a hypothesized mechanistic relationship.

### vtaVNS reduces distance to criticality

The association between reduced visual ERP and successful memory encoding suggests that vtaVNS may promote efficient allocation of cortical resources, such that reduced neural computation for visual processing leaves greater capacity for working-memory maintenance and updating. A near-critical operating regime enables enhanced tunability in response to behavioral demands, providing a potential network mechanism for this redistribution^49^. We next tested the third prediction of the cortical-efficiency hypothesis that vtaVNS shifts cortical dynamics toward criticality. Our multivariate linear model analysis showed that the distance to criticality is smaller under active vtaVNS than under sham (**Fig. 3C, Fig. S4C**, Frobenius norm of the electrode-wise coefficient vector = 0.073, permutation p = 0.003, permutation test, see **Supplementary Materials** for details). The reduction was largest over parietal electrodes (*β*_*P*4_ = −0.032, standardized effect = −0.470 residual SD for P4; *β*_*P*3_ = −0.020, standardized effect = −0.312 for P3). Although vtaVNS reduced the distance to criticality, trial-by-trial variations in distance to criticality were not associated with successful encoding (p = 0.996, permutation test). This suggests that vtaVNS shifts cortical dynamics toward a more critical operating state under which trial-specific neural processes, such as cortical response to visual stimuli, influence the probability of memory encoding. Based on these findings, we proposed a mechanistic model explaining the improvement of working memory via vtaVNS. Specifically, vtaVNS engages the LC-NE pathway, shifting cortical dynamics toward a near-critical operating regime, within which cognitive resources are flexibly allocated and efficiently utilized among perception, memory updating, memory maintenance, and executive control to meet behaviorally relevant demand **(Fig. 3D)**.

### Exploratory analysis of heartbeat-evoked activity

N-back tasks require dynamic allocation of cognitive resources between encoding of visual stimuli and updating and maintaining task-relevant representations^50^. During the task, participants were not instructed to monitor their heart rate, nor was cardiac information required for task performance. If vtaVNS promotes cortical efficiency, we expected it to reduce the neural activity required for interoceptive processing, in addition to sensory processing. Cardiac activity is a canonical interoceptive signal whose cortical processing has been measured using heartbeat-evoked potentials (HEPs)^44,51,52^. We therefore explored whether vtaVNS reduced HEP during the N-back task.

HEP waveform consistency was only slightly greater after the heartbeat than during the pre-R-peak baseline for a subset of electrodes and was not reliably elevated across channels (**Fig. S2A, B**, median under the ROC curve based on inter-trial correlation = 0.510). Cluster-based permutation tests with TFCE revealed no significant effects of vtaVNS on HEPs within the neural component (**Fig. S2C**). We considered two explanations. First, vtaVNS may have little to no effect on interoceptive processing in a working memory task. Second, our component-separation procedure may classify some HEP signal as part of the cardiac component if the cortical response to the heartbeat occurs with a relatively consistent latency and amplitude relative to the QRS complex. To test the second possibility, we repeated the cluster-based permutation tests on the cardiac component and found clusters of reduced cardiac component amplitude in the vtaVNS condition compared to sham (**Fig. S2D)**. The largest spatiotemporal cluster was frontocentral, centered at the F4 electrode, and extended from 0.404 to 0.672 s after the R-peak (permutation test, p < 0.001, largest difference = -0.308 z-score unit). The location and timing align with a previous report of HEP in humans^51^, but the timing was earlier than a previous study reporting the effect of vtaVNS on HEPs^52^. To confirm that the change in the cardiac component was not due to a change in the ECG signal, we compared ECG signals time-locked to the heartbeat and found no difference (**Fig. S2D**, permutation test). We interpret these results as tentative evidence that vtaVNS reduced cortical dynamics implicated in interoception.

## Discussion

Across scalp EEG and intracranial LFP recordings, our findings indicate that 6 Hz vtaVNS promotes cortical efficiency during working memory. Specifically, vtaVNS reduced cortical responses to visual stimuli during the N-back working memory task, and smaller visual ERPs were associated with successful encoding in the 4-back task. In parallel, vtaVNS shifted cortical dynamics toward criticality, the boundary between subcritical and supercritical regimes. These observations are difficult to reconcile with the speculated sensory-amplification hypothesis or with an alternative explanation, generalized cortical suppression, because both predict that brain dynamics move away from criticality rather than toward it. Sensory amplification would drive the network toward supercritical dynamics, whereas suppression would drive it toward subcritical dynamics. Together, our findings provide a physiological link between the engagement of the LC-NE pathway via vtaVNS and improvements in working memory.

Physiological evidence from our group and independent research groups demonstrates that vtaVNS activates LC activity and modulates the LC-NE system^14,53,54^. In the present study, both visual ERP and intracranial high-gamma activity reflect large-scale synchronous neuronal activity^42^. The reduced visual ERP and high-gamma activity in our study align with previous findings that NE is associated with desynchronized cortical activity^55^. However, physiological consequences of LC-NE modulation via vtaVNS remain poorly understood, particularly in the context of working memory. The adaptive gain framework proposes that phasic LC activity selectively increases neural sensitivity around behaviorally relevant events, whereas tonic LC firing produces a relatively uniform amplification of the neuronal response, rendering the system more sensitive to task-irrelevant stimuli^26,27,56^. The Glutamate Amplifies Noradrenergic Effects (GANE) provides a biological account through which the LC–NE system regulates cortical efficiency. Neurons actively engaged in behaviorally relevant processing generate strong glutamatergic activity, which can promote local NE release and create NE “hot spots”^57^. Within these hot spots, NE engages low-affinity β-adrenergic receptors, which enhance neuronal excitability and presynaptic glutamate release^57,58^. Elsewhere, NE remains at lower concentrations and preferentially engages high-affinity inhibitory adrenergic receptors, relatively suppressing background neural activity. Thus, one interpretation of our reduced cortical responses to visual stimuli is that stimulus-paired vtaVNS increases the selectivity of neural processing, allowing task-relevant representations to be processed with less neural recruitment. Our finding that vtaVNS reduced distance to criticality represents a systems-level consequence of LC–NE modulation. Recent animal studies provide emerging evidence linking LC activity to critical brain dynamics^59,60^. Together, it is possible that vtaVNS-mediated LC-NE modulation shifts cortical networks toward a near-critical regime that supports efficient information processing^29^.

However, trial-by-trial variation in visual ERP amplitude was associated with successful memory encoding, whereas variation in distance to criticality did not. This dissociation suggests that the near-critical regime induced by vtaVNS is a global neuromodulatory state that supports working memory rather than a direct neural marker of memory. This operating state provides the computational conditions under which trial-specific neural processes, as reflected in visual ERPs, influence memory encoding. Under this interpretation, the modulation of the LC–NE system via vtaVNS shifts cortical dynamics toward a more critical operating state, in which neural resources may be allocated more selectively and flexibly, thereby increasing the capacity to update and maintain working memory^12,14,38,61,62^.

Our mechanistic model of vtaVNS generates two predictions about its behavioral effect: vtaVNS is more beneficial when distracting sensory input competes with mnemonic processing for finite cortical representational capacity. Conversely, simple detection of changes may not benefit from vtaVNS. Indeed, a study reported no behavioral effect of vtaVNS during change-detection tasks^54^. Moreover, our findings that reduced visual ERPs were associated only with successful encoding in the 4-back task imply that the benefit of cortical efficiency is maximized for working memory tasks when cognitive resources are at the edge of shortage. A dual behavioral task could directly test whether our prediction is true. Specifically, researchers could combine the N-back task with infrequent external probes in which participants maintain working-memory items while occasionally detecting a subtle luminance change. Under our mechanistic model, vtaVNS is expected to improve memory but not probe detection.

Our study has limitations. First, the pathology of patients with epilepsy limits the generalizability of our high-gamma finding. To ensure feasibility in the neurosurgical setting, we adapted the working-memory paradigm to reduce task burden and used within-block randomization of stimulation and no-stimulation trials, providing a strong control that minimizes confounding by block-level temporal correlations. This design was optimized to confirm the physiological effects of vtaVNS but not to determine the duration of the behavioral effects of vtaVNS. Second, our exploratory analysis of HEPs showed attenuated cortical responses to cardiac signals, supporting that vtaVNS may attenuate processing that is less relevant to the current behavioral goal. Given that the physiological characterization of HEP is under active investigation, accurate interpretation of HEP findings requires better physiological understanding of HEP and biophysical modeling of the pathway. Third, the earlobe stimulation used as the sham condition may have been physiologically active^63^, and the concha and earlobe may differ in mechanical coupling, producing unequal somatosensory or auditory input. The active-versus-sham differences observed here support a location-dependent effect, but they are not definitive evidence of vagal-pathway specificity. A study reported that tail stimulation in rodents can alter LC activity, suggesting that several sensory pathways may act on the LC–NE system^64^. Biophysical studies of vagus nerve stimulation could consider a sensation-matched control delivered at a distant site without vagal innervation to test whether auricular vtaVNS recruits the LC more strongly. Resolving this limitation may reveal an opportunity to control LC–norepinephrine activity through spatially and temporally patterned stimulation across multiple sensory pathways. Fourth, although the observed reduction in distance in active vtaVNS conditions supports flexible computation, our data do not directly demonstrate that cognitive resources are reallocated toward mnemonic processing. The distributed nature of working memory makes it difficult to capture neural activity associated with memory maintenance using electrophysiology^65^. Scalp EEG provides limited functional specificity, whereas intracranial recordings provide greater specificity but sample part of the mnemonic network. Advances in brain-sensing technology and computational modeling may help capture distributed mnemonic representations and determine whether vtaVNS enables neural resources to be flexibly reallocated in response to behavioral demands. Lastly, our proposed mechanism of action for vtaVNS is based on 6 Hz stimulation. Future studies should examine other commonly used taVNS frequencies to determine their effects on working memory and whether our proposed mechanism of action generalizes across frequencies^9,41^.

In summary, our data argue against a model in which vtaVNS improves working memory by uniformly amplifying cortical responses to external stimuli. Instead, vtaVNS improves cortical efficiency, allowing cognitive resources to be flexibly reallocated to task-relevant mnemonic processing. Our model predicts the greatest benefit when exteroceptive and mnemonic demands are in direct competition.

## Data and code availability

All analysis scripts and data for statistical analysis and figure plotting are available on a GitHub repository (https://github.com/GanshengT/taVNS-working-memory-ephys). Additional data, such as imaging data from patients with epilepsy and the source files for 3D printing the custom vtaVNS device, are available from the corresponding authors upon reasonable request. Intracranial electrode localization source code and relevant data are available on a GitHub repository (https://github.com/GanshengT/intracranial_contact_loc).

## Acknowledgements

This work was supported by NIH/NIMH (R01-MH120194), NIH/NIBIB (P41-EB018783, R01-EB026439), McDonnell Center for Systems Neuroscience, Fondazione Neurone, NIH/NINDS (U24-NS109103, U01-NS108916, U01-NS128612, R21-NS128307), and NIH/NEI (R01-EY037195). We acknowledge insightful discussions regarding visualization and computational modeling from Dr. Sai Sun and Noah Bryson.

## Author contributions

Conceptualization: G.T. and E.C.L.; Data curation: G.T., P.B., E.C.L., J.R., and J.T.W.; Formal analysis: G.T.; Funding acquisition: E.C.L. J.L.G., and P.B.; Investigation: G.T., P.B., and E.C.L.; Methodology: G.T., P.D., and T.K.; Software: G.T. and P.B.; Supervision: P.B. and E.C.L.; Validation: G.T., P.B., T.K., and E.C.L.; Device: J.A. and J.L.G.; Visualization: G.T.; Writing – original draft: G.T.; Writing – review & editing: G.T., and E.C.L.

## Declaration of generative AI and AI-assisted technologies in the writing process

During the preparation of this work, the author(s) used ChatGPT 5.5 and Grammarly AI to improve the readability of the manuscript. After using these tools, the authors reviewed and edited the content as needed and take full responsibility for the content of the published article.

## Methods

### Subjects

20 healthy individuals aged 18 to 45 years, without significant past medical history and with no known neurological deficits, completed this study. Participants were required to abstain from alcohol and caffeine consumption starting one day before the study. We also recruited 20 patients with intractable epilepsy who were undergoing invasive monitoring for localizing the seizure onset zone. The study was performed in accordance with the Declaration of Helsinki, the International Conference on Harmonization Good Clinical Practice Guidelines, and the applicable United States Code of Federal Regulations. The study was approved by the Washington University in St. Louis Institutional Review Board (IRB ID#: 202101161). Written informed consent was obtained from all study participants.

### Vibrotactile taVNS (vtaVNS)

The vtaVNS system comprises parts fabricated with stereolithographic additive manufacturing (**Fig. 1C**). The system includes an ear cup, a cushioning base that provides soft contact with the head, and a T-head bolt that connects the contact tip and the ear cup via ear cup covers. An eccentric rotating mass (ERM) vibration motor of 5 mm diameter (Vybronics Inc, Brooklyn, NY) is held in at the contact tip^14^. The motor’s power cable passes through the T-head bolt. The T-head bolt allows individual adjustment of the contact depth. In addition, rotating the T-head bolt allows switching the stimulation target between the concha area and the earlobe. A digital stimulation box (g.STIMBOX, g.tec medical engineering, Austria) supplied 5V power at 6 Hz to the eccentric Rotating Mass vibration motor. An elastic strap stabilizes the vibrotactile device on the head.

To improve wearability and comfort for patients with implanted electrodes, we redesigned the vtaVNS device as a compact, 3D-printed ear clip fabricated from Flexible 80A Resin (Formlabs). The clip has an A-shaped configuration formed as a single printed structure. Two opposing arms cross at a central cylindrical region. One arm terminates in a rounded contact end housing the ERM vibration motor, whereas the opposing arm terminates in a flat, flexible contact surface with a central opening for stable positioning on the ear. Both arms slightly extend beyond the central cylinder to form handles. The elastic force of the printed resin keeps the two contact ends closed, and squeezing the handles together opens them for placement on the ear.

### Study design

Healthy participants completed N-back tasks. Subjects responded with a button press when they thought the currently presented digit was the same as the digit presented N trials prior. The digits presented in the N-back task were randomly selected from 4 digits: 0, 1, 2, and 3. Participants completed three sessions of N-back tasks: a baseline session, a sham session, and a VNS session, with the order of sessions randomized. To mitigate potential confounding factors related to fatigue or residual stimulation effects, participants rested for 3 min between sessions. During the baseline session, the participant completed 8 blocks of N-back tasks with N ranging from 1 to 8. Between two blocks, subjects were given a mandatory minimum rest period of 10 s, after which they could proceed to the next block by pressing a button. Each block included 40 probes, that is, digit presentations, when the subject needed to decide whether to press the button. Before the first probe, N digits were presented to load the participant’s working memory. Throughout the task, digits were displayed in white against a black screen background for 1 s each. Between the presentation of two digits, a red fixation cross was displayed in red for 0.5 s. During the active vtaVNS or sham session, the participant completed 8 blocks of N-back tasks with N ranging from 0 to 4. The 0-back task in the active vtaVNS or sham session was designed as a control for attention.

The active vtaVNS, sham control, and baseline session only differed in the stimulation site. In active vtaVNS session, vibrotactile stimulation was delivered to the left cymba concha area of the outer ear; and in sham session, vibrotactile stimulation was delivered to the left earlobe. In both active vtaVNS and sham sessions, stimulation was paired with the presentation of the digits. In other words, vibrotactile stimulation was on while the digit was displayed and deactivated during the presentation of the fixation cross. This protocol was designed to reduce cutaneous mechanoreceptor adaptation. During the baseline session, subjects wore the vtaVNS device that contacted the earlobe. Before the study, the participants received instruction from the experimenter and practiced the N-back task until they were familiar with it.

To assess cortical response to vtaVNS, we recorded local field potentials from patients with epilepsy during spatial working memory tasks. During pilot testing, N-back task used in the healthy population was poorly tolerated in patients with epilepsy due to postoperative fatigue, epilepsy-related burden, task difficulty, and the rapid presentation of digit stimuli, which reduced engagement and limited reliable task performance. We therefore adopted a classic spatial working memory paradigm that is less demanding and well tolerated, while measuring the effect of vtaVNS on cortical responses to visual stimuli during memory encoding.

Thus, we adopted a classic spatial working memory task aiming to study the effect of vtaVNS on cortical responses to visual stimuli during memory encoding. After practice, participants completed multiple sessions of the spatial working memory task. In active vtaVNS session, the vtaVNS device (ear clip) was placed on the cymba concha; and in sham session, the same device was placed on the left earlobe. The number of sessions was determined by the clinical schedule and participant availability. Each session consisted of 10 trials, which were randomized to receive either vibrotactile stimulation or no stimulation. Each trial began with the presentation of an empty grid, followed by the appearance of a red cross, which cues participants to the center of the dot pattern during the subsequent encoding phase. During the retrieval phase, participants used a keyboard to indicate whether the dot’s location matched that of any dot shown during encoding. To minimize fatigue and vtaVNS habituation, trials started after the subject pressed a key, with a mandatory 6-second inter-trial interval.

### Data acquisition

During the N-back task in healthy participants, we simultaneously recorded EEG and ECG using a g.USBamp amplifier (g.tec medical engineering, Austria). We recorded EEG at 1,200 Hz from 16 scalp locations using g.SAHARA active dry electrodes (g.tec medical engineering) positioned according to the international 10–20 system, providing coverage of frontal, temporal, parietal, and occipital regions. Each g.SAHARA electrode has an integrated amplifier and consists of eight gold-alloy contact pins designed to maintain low electrode–skin impedance. Before recording, we adjusted the cap position and individual electrode contact as needed, guided by electrode–skin impedance measurements obtained using the g.USBamp impedance-measurement mode. We also visually inspected the continuous EEG to identify excessive noise or unstable signals and made further adjustments as needed. We recorded ECG using adhesive foam electrodes placed bilaterally on the upper chest below the clavicles and on the left and right ankles. We connected the left ankle electrode to both the reference and ground inputs of the amplifier and connected the right ankle electrode to the equipotential grounding lead. We derived ECG from the differential voltage between electrodes positioned bilaterally on the upper chest. We recorded gaze position and pupil diameter at 60 Hz using a Tobii Pro Fusion eye tracker (Tobii, Sweden). We used BCI2000 to coordinate stimulus presentation, behavioral recording, physiological data acquisition, and event synchronization, including photodiode signals and vibrotactile stimulation triggers. To measure visual-stimulus onset precisely, we attached a photodiode (g.tec medical engineering GmbH, Austria) to the bottom-left corner of the display. At each digit onset, a white marker appeared at the corresponding screen location, which was detected by the photodiode.

To investigate the effect of vtaVNS on population neural response to visual stimuli, we recorded intracranial EEG at the bedside in the epilepsy monitoring unit using a clinical SEEG system (Neurofax EEG-1200, Nihon Kohden, Irvine, CA) equipped with a 256-channel JE-120 amplifier. We sampled signals from depth electrode contacts at 2,000 Hz with 16-bit resolution (±3.2 mV input range, 97.65 nV/LSB). Per standard clinical practice, two subgaleal electrodes served as the reference and ground during signal acquisition. We used a desktop computer running BCI2000 to synchronize stimulus presentation, behavioral responses, and electrophysiological recordings. We presented stimuli on a 27-inch monitor (TUF Gaming VG27AQL1A, ASUS; 2560 × 1440-pixel resolution) mounted on a mobile cart with an adjustable monitor arm, allowing us to adjust the display position to each participant’s posture and bed orientation.

### Preprocessing of scalp EEG data

To characterize neural responses to visual stimuli, we processed continuous EEG data to derive visual event-related potentials (ERPs) while separating artifactual components including blink-related signals, stimulation artifacts, and cardiac artifacts. We first removed line noise and slow drifts, identified periods of unreliable recording, and then estimated blink-, vibrotactile-, and cardiac-related components using a template-matching algorithm.

To remove line noise at 60 and 120 Hz, we applied Zapline-plus, a method that combines spectral and spatial filtering^66^. We then estimated potential slow changes in skin potentials recorded by EEG using empirical mode decomposition (EMD) and subtracted the estimated trend from the EEG data. Transient disruptions in electrode–scalp contact, for example during head movement, can produce abnormally large and abrupt voltage changes, or flat signals. As a quality control, we therefore identified periods of unreliable recording by comparing signal amplitude and its temporal derivative to robust z-scores based on the median and median absolute deviation (MAD). We required detected abnormalities to persist for at least 10 ms. We extended the detected period by 50 ms on either side to capture the full contaminated signal. These criteria flagged 1.6% of EEG data as unreliable. We did not exclude these detected periods during supervised sequential artifact-component separation. When extracting visual ERPs, we excluded stimulus-presentation-locked epochs for each electrode that overlapped the detected periods of unreliable recording.

### Identification of heartbeat timing

Separating cardiac-artifact components from EEG signals requires Heartbeat timing to construct the template used. We defined the timing of each heartbeat by the R-wave peak. To improve the signal-to-noise ratio of the ECG waveform for reliable R-wave delineation, we bandpass-filtered the ECG from 1 to 40 Hz using a third-order Butterworth filter in both the forward and reverse directions. We removed the constant-voltage offset by subtracting the center of the most frequently occupied bin from an ECG amplitude histogram. We then delineated the ECG waveform using *ECGdeli*, a toolbox that detects the onset, peak, and offset of the P wave, QRS complex, and T wave^67^.

### Identification of blink timing

To obtain a reliable set of seed blink events for constructing the initial EEG blink-artifact template, we identified blink onsets and offsets from pupillometry, and cross-validated them against electrooculogram (EOG). We identified candidate blink onsets and offsets when constriction or dilation velocity deviated from the mean by more than 5 standard deviations. Because recorded pupil diameter has finite measurement resolution, we smoothed pupil diameter using a 10-ms moving average before calculating constriction or dilation velocity. We paired detected blink onsets and offsets according to the expected blink duration (∼250 ms). We retained physiologically plausible blink events with durations ranging from 50 ms to 1 s.

We cross-validated pupil-derived blink events using EOG. We derived the EOG by subtracting the mean signal across posterior EEG electrodes from the mean signal across two frontopolar electrodes (AF3 and AF4). To quantify blink-related activity captured by the EOG, we bandpass-filtered the EOG signals from 0.5 to 8 Hz using a second-order zero-phase Butterworth filter and calculated their signal energy. We identified blink events using a threshold-based criterion (median-based z-score of signal energy > 16). Eye-tracking data were unavailable for three participants due to logistical constraints; for these participants, we identified blink events directly from the EOG.

Temporal alignment of detected blink onsets was important for constructing a blink-artifact template because the detected blink onset times may not correspond to the onset of the blink-related EEG artifact. Within each subject, across blinks, EEG waveforms at frontal electrodes following a blink are expected to be stereotyped. Thus, we aligned blink onsets within a temporal window (±120 ms) by shifting the corresponding frontal EEG waveform to maximize similarity among their temporal derivatives. We subsequently used the aligned blink onsets to construct the blink-artifact template.

### Supervised sequential artifact-component separation

Blinks, vibrotactile stimulation, and cardiac activity produced stereotyped voltage patterns that could overlap with the visual ERPs. To separate these three artifact components from the EEG signals, we first construct component templates based on signals around the event onset and construct artifact components by template matching. We then sequentially removed them:

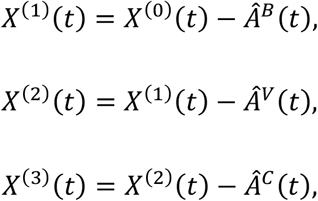

where 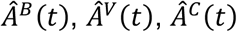 denote the estimated artifactual components related to blink, vibrotactile stimulation, and cardiac activity, respectively; *X*^(2)^(*t*) is the neural component.

For an event *e* such as a blink, let **T**(*u*) denote the location-specific artifact template as a function of relative time (u), defined over a finite interval *W*. We define **T**(*u*) = 0 for *u* ∉ *W*. The fitted artifact has the form 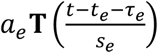, where *t*_*e*_ is the event onset (e.g., blink onset), *τ*_*e*_ is an event-specific temporal shift, *s*_*e*_ is an event-specific temporal stretch, *a*_*e*_ is the amplitude scale. Thus, each event contributes a shifted and scaled copy of the finite-duration template to the continuous EEG. The artifact component 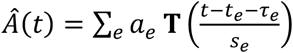

### Blink component estimation

We estimated the blink-artifact component using a two-stage procedure: template-based detection of blink events, followed by reconstruction of their multichannel EEG waveforms. Blink times identified via pupillometry or EOG served as seed events to construct an initial blink-artifact template, which we used to identify all blinks throughout the recording. To detect blinks, we applied template matching to spatially whitened EEG data, as whitening can reduce the influence of electrode-level covariance differences on multichannel matching. Whitening is used only for blink detection. We later estimated the templates for artifact-component construction based on the original, unwhitened EEG signals.

To construct the initial template, we extracted EEG signals from −0.15 to 0.30 s around each pupil- or EOG-derived blink onsets. We subtracted the mean amplitude of the baseline interval (−0.15 to −0.05 s). We constructed the initial template by averaging blink events after excluding the most extreme 5% of values at each end (i.e., trimmed mean), to preserve the typical blink-related waveform. The initial template is an electrode-by-time matrix. For each electrode, we determined a finite interval over which the template would contribute to the reconstructed blink-artifact component. We searched progressively wider intervals within the extracted segment, using 2-ms steps. We selected the first interval whose endpoint amplitudes differed by no more than 10% of that electrode’s peak-to-peak amplitude. Within the selected interval, we subtracted the mean of the two endpoint amplitudes, making both endpoints close to zero, and set the template to zero outside that interval. These operations were also applied to the stimulation-artifact and cardiac-activity templates to reduce discontinuities at the boundaries of the finite-duration templates.

To detect blinks throughout the recording, we slid the initial template across the whitened EEG. For each time point, we evaluated both the waveform similarity of the EEG signals to the template and their amplitude agreement with the template. For location *l*, let *y*_*l*_ denote the signal vector and *T*_*l*_ denote the template vector. We quantified waveform similarity using cosine similarity 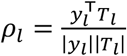. We quantified amplitude agreement using a normalized goodness-of-fit score 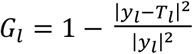. We combined these two metrics across locations into a composite score for template Matching 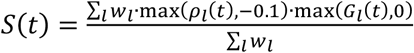, where *w* denotes the location-specific weight. This composite score favored time points that exhibited a blink-like waveform and matched the expected blink-artifact amplitude.

The location-specific weight *w*_*l*_, defined in Eq. (**Error! Reference source not found**.), emphasizes locations with a clear blink waveform. We assigned zero weight to electrodes whose template energy *E*_*l*_ = ∑_τ_ *T*_*l*_ (*τ*)^2^ was no greater than 1% of the mean energy. We then quantified the prominence of the blink waveform 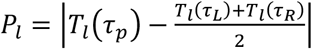, where *τ*_*p*_ is the time of the most prominent peak of template energy, and *τ*_*L*_ and *τ*_*R*_ are the associated left and right flanking minima. We defined the location-specific weight as linearly scaled prominence,

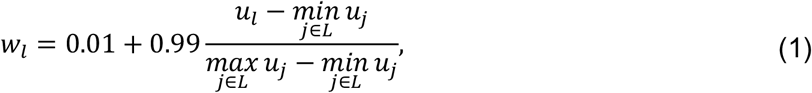

Where 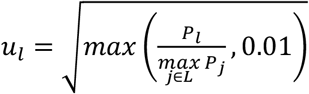.

To identify blink onsets from the continuous template-matching scores, we derived a time-varying detection threshold from surrogate templates that preserve signal characteristics while disrupting the spatiotemporal pattern. We generated 10 surrogate templates using Fourier Phase Randomization and shuffling temporal blocks within electrodes. At each time point, we defined a detection threshold as the 99th percentile of the corresponding surrogate matching scores. We identified candidate blink intervals in which the actual matching score exceeded this threshold by more than one standard deviation and remained above this threshold for at least 15% of the template duration. For each identified interval, we defined a blink time as the point of greatest exceedance above the time-varying surrogate threshold. We combined two identified intervals if their blink time was separated by less than 250 ms. We used these identified blink times to construct the final template for extracting the blink component from the unwhitened EEG signals, following the same procedure applied to derive the initial template.

Because individual blink artifacts can vary in timing, duration, and amplitude, we fitted the final blink template to each detected blink in the EEG signals: 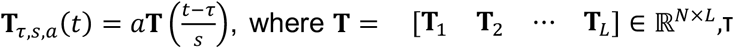 is a temporal shift parameter, *s* is a duration-scaling parameter, and *a* is an amplitude-scaling parameter *a*. We removed local linear trends from both the EEG signals around the detected blink and the template to prevent slow baseline changes from influencing the fit. We estimated parameters τ, *s* by searching for a combination that minimized the weighted proportional signal energy,

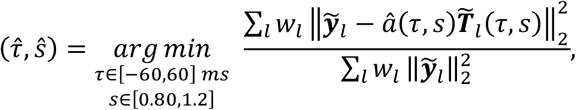

where 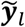 is the EEG segment with local linear trend removed for location 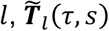 is the template after temporal shifting and duration scaling, and *w*_(_ is defined in Eq. **Error! Reference source not found**.. For each candidate (*τ, s*), the least-squares solution for the amplitude-scaling factor that minimizes weighted proportional signal energy 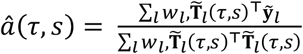. We required *â*∈(0.5,4).

To avoid poorly matched templates, we accepted an event-specific blink component only when its goodness-of-fit score 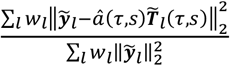 was finite and exceeded the 10th percentile of goodness-of-fit score obtained by applying the same fitting procedure to the seed blink events used to construct the initial template. This empirical threshold required newly detected blinks to achieve a fit quality comparable to the cross-validated blink events.

### Vibrotactile stimulation artifact component estimation

We estimated the vibrotactile stimulation artifact from the residual EEG signals after removing the blink component. We derived stimulation timing from electrical pulses delivered to the vibrotactile motor, recorded via a trigger box (g.TRIGbox, g.tec medical engineering GmbH, Austria). Because all stimulation onsets were recorded, we did not need to detect the full stimulation event using a seed template, unlike in blink component estimation. For each electrode, we extracted EEG epochs aligned to the onset of each stimulation and calculated a stimulation-locked template using a 10% trimmed average. Each 6-Hz stimulation train consisted of repeated vibrotactile pulses, such that the stimulation-locked average contained both brief, highly stereotyped pulse-related artifacts and physiological activity that varied more across trials. We therefore separated them by first identifying time intervals dominated by the stereotyped artifact and then using the remaining intervals to estimate the physiological signal. To identify time intervals dominated by the pulse-related artifact, we exploited its expected sharp and stereotypical waveform across stimulation trials. Specifically, for each electrode, we quantified waveform consistency across stimulation trials using Pearson correlation within 20-ms moving windows. We then averaged the consistency across electrodes and defined stimulation-artifact intervals as periods in which the mean consistency exceeded the 80th percentile of its values across time. We expanded each identified interval by 10 ms on either side. To derive physiological signals, we fitted a sixth-order Fourier series to the stimulation-locked average outside the artifact intervals:

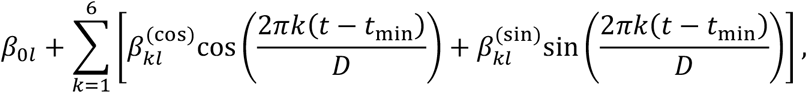

where *D* = *t*_max_ − *t*_min_ is the duration of a stimulation train. We defined the stimulation-artifact template as the residual obtained by subtracting the fitted physiological signals from the stimulation-locked average. We fitted the template to the residual EEG signals after removing the blink component following the procedure described in **Methods: Blink component estimation**. For baseline sessions, in which no vibrotactile stimulation was delivered, the stimulation-artifact component was defined as zero.

### Cardiac component estimation

We estimated the cardiac component from the residual after subtracting the blink and, where applicable, stimulation-artifact components. We extracted EEG epochs centered on R peaks detected from the ECG (see **Methods: Identification of heartbeat timing**), baseline-demeaned epochs using a pre-R-peak interval (−0.30 to −0.10 s), and constructed a heartbeat-locked template using a 10% trimmed average. Because the cardiac artifact exhibited a simple, stereotyped waveform, we fitted the template to each EEG segment (−0.10 to 0.40 s), using parameters *τ* and *a* representing temporal shift and amplitude scaling, respectively, to account for beat-to-beat variation, and an intercept parameter *b* to account for the local voltage offset.

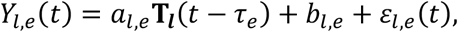

where **T**_***l***_ is the heartbeat-locked template at location *l* and *e* denotes the heartbeat event. For a given shift *τ*_*e*_, we estimated the amplitude and local voltage offset using a least-squares solution. We searched for *τ*_*e*_ that minimized the total template-fitting error:

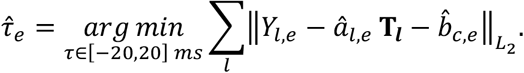

We evaluated goodness of fit for each heartbeat relative to a linear model

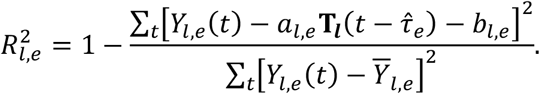

We constructed the cardiac component as

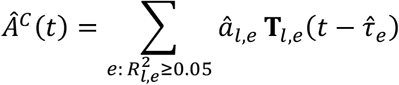

### Visual event-related potential (ERP) waveform consistency

Visual ERPs are expected to have similar waveforms following stimulus presentation. To verify that the neural component in EEG signals retained this characteristic response, we quantified waveform consistency across stimulus presentations within stimulus-locked epochs (-1.0 to 1.5 s relative to stimulus onset). Because scalp EEG is susceptible to electrode–scalp contact disturbances, especially during movement, it may contain abrupt voltage transients that were not captured by the preceding artifact-component separation procedure. To identify epochs containing such transients, we calculated the rate of change of voltage over a 50-ms lag throughout each epoch and summarized it by its root-mean-square value. We identified epochs with artifactual transients using a threshold defined as the median plus four scaled median absolute deviations. For each electrode, we quantified intertrial waveform consistency by correlating each retained epoch with the average of all other retained epochs. To determine whether waveform consistency increased following stimulus presentation relative to baseline, we compared the distributions of post-stimulus (0 to 0.5 s) and baseline (-0.5 to 0 s) Pearson’s correlation values using a receiver operating characteristic (ROC) curve. Across correlation thresholds, we defined the true-positive rate as the fraction of post-stimulus correlations exceeding the threshold, and the false-positive rate as the fraction of baseline correlations exceeding the same threshold. The area under the ROC curve (AUC) quantified their separability, with an AUC greater than 0.5 indicating greater waveform consistency following stimulus presentation than during baseline^68^.

### Heartbeat-evoked potential (HEP) waveform consistency

To determine whether the heartbeat-locked activity exhibited consistent waveform, we quantified waveform consistency within epochs spanning −2.0 to 2.0 s relative to each R peak. For each electrode, we quantified waveform consistency using the same leave-one-out Pearson correlation procedure described in Methods: Visual event-related potential (ERP) waveform consistency. We compared correlation values during the post-R-peak interval (0 to 1.0 s) with those during the preceding baseline interval (−1.4 to −0.4 s) using area under the ROC curve analysis.

To determine whether visual-ERP and HEP waveforms were more consistent following stimulus presentation and R peaks, respectively, than during the preceding baseline, we modeled AUC values using linear mixed-effects models. For each electrode, we fitted a linear mixed-effects model, with *AUC* − 0.5 as the dependent variable and stimulation condition (baseline, sham, and active vtaVNS) as a fixed effect. We included subject as a random intercept. The model intercept therefore estimated the mean AUC relative to chance across baseline, sham, and vtaVNS conditions. We tested whether the intercept was greater than zero, corresponding to an AUC greater than 0.5. We applied Benjamini–Hochberg correction across the 16 EEG electrodes.

### Assessing effects of vtaVNS on visual ERP

Based on the cortical-efficiency hypothesis, we predicted reduced visual ERP amplitudes during active vtaVNS. To test this hypothesis, we calculated visual ERPs as the arithmetic mean of EEG epochs time-locked to visual stimulus presentations, separately for each subject, electrode, task-difficulty level, and condition (vtaVNS, sham, and baseline). To prevent differences in prestimulus voltage offsets from contributing to ERP amplitude differences across difficulty levels and conditions, we subtracted mean visual ERP amplitude during the 500 ms preceding stimulus onset from visual ERPs. Baseline ERPs for one participant at levels N = 1–4 overlap the identified unreliable recording. We imputed these waveforms using the corresponding means from the other 19 subjects, adjusted by this participant’s average deviation from the means of other participants in the sham and vtaVNS conditions at the same difficulty level.

Visual ERPs are considered physiological measures of visual attention and sensory processing^69^. To clarify whether visual ERPs could also reflect processes associated with working memory updating or maintenance, we examined the relationship between baseline visual ERP amplitude and working memory load (N = 1–8). Let *A*_*jnl*_(*t*) denote the visual ERP amplitude for participant *j*, working memory load *n*, location *l*, and time *t*. Because we tested within-participant differences across working memory loads, we standardized visual ERP amplitude separately for each participant and electrode using a single mean and standard deviation pooled across all difficulty levels and epoch time points:

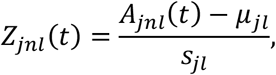

where *A*_*jnl*_(*t*) is the mean visual ERP amplitude within the 50-ms window centered at *t*. Window centers were advanced in 25-ms steps. We fitted an ordinary least squares regression to the mean visual ERP amplitude across working memory loads: *Z*_*jnl*_(*t*) *∽ α*_*jl*_(*t*) + *β*_*jl*_(*t*)*N* + *ε*_*jnl*_(*t*). We standardized estimated *β*_*jl*_(*t*) relative to the subject’s residual variability:

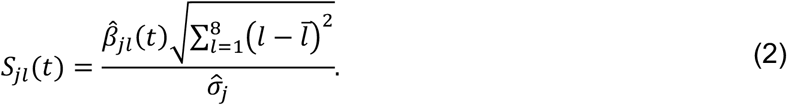

We estimated the subject’s residual variability 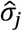 using median-based variability estimation: 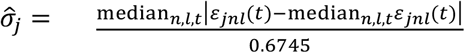. We assessed the statistical significance of the association between visual ERP amplitude and working memory load using a two-sided, one-sample permutation test with threshold-free cluster enhancement (TFCE). See **Cluster-based permutation tests with threshold-free cluster enhancement (TFCE)** for details.

To determine the effect of vtaVNS on visual ERP amplitudes, we fitted a linear mixed-effects model separately for each location and time:

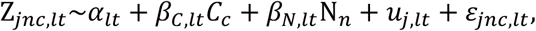

where Z_*jncl*_(*t*) denotes standardized visual ERP amplitudes within the 100-ms window centered at *t* for participant *j*, difficulty level *n*, condition *c*, location *l* ; *α*_*lt*_ denotes the intercept; *β*_*C,lt*_ represents the fixed effect of stimulation condition; *β*_*N,lt*_ represents the fixed effect of working memory load; *u*_*j,lt*_ is a participant-specific random intercept; and *ε*_*jnclt*_ is the residual error. We coded stimulation condition as *C*_*c*_ = −0.5 for sham and *C*_*c*_ = +0.5 for vtaVNS. Thus, *β*_*C,lt*_ estimated the vtaVNS–sham difference in visual ERP amplitude after accounting for working memory load. We standardized *β*_*C,lt*_, following Eq. (**2**), and assessed their statistical significance using a two-sided permutation test with TFCE.

### Cluster-based permutation tests with threshold-free cluster enhancement (TFCE)

Testing the effects of vtaVNS on visual ERP amplitude across many electrodes and time creates a multiple-comparison problem. Cluster-based permutation testing controls the family-wise error rate by comparing statistics from spatially and temporally contiguous clusters against a permutation-based null distribution. However, defining these clusters requires a prespecified statistical threshold. To avoid reliance on a single cluster-forming threshold, we used threshold-free cluster enhancement (TFCE), which enhances statistics based on their magnitude and the extent of contiguous effects across spatially adjacent electrodes and consecutive temporal windows^46^. Cluster-based permutation tests with TFCE are well suited for assessing the effect of vtaVNS on visual ERP amplitudes because physiological effects of vtaVNS are thought to persist across location and time.

To apply TFCE to a location-by-time statistic map *T*, at each threshold magnitude *h*, we transformed each statistic into an enhanced score that incorporated both the magnitude of the statistic and the extent of the surrounding same-sign effect. As an example, to apply TFCE on *S* in Eq. (**2**), the standardized slope was converted into a T-like statistics, 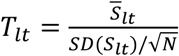, where *N* is sample size. At each threshold magnitude *h*, we identified connected sets of statistics that exceeded *h* or were below −*h* for negative effects. We defined connectivity across spatially adjacent electrodes and consecutive temporal windows.

For each location *l* and time window *t*,

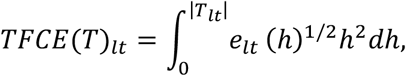

where *e*_*lt*_(*h*) is the number of statistics in the same-sign connected component containing statistic at location *l* and time window *t* at threshold *h*. Thus, *TFCE*(*T*)_*lt*_ is larger when *T*_*lt*_ was stronger and belonged to a more extensive contiguous effect. Positive and negative effects were evaluated separately. Programmatically, thresholds started at zero and advanced in increments of 0.1. Spatial adjacency was derived from the standard 10–20 EEG electrode montage.

We assessed statistical significance using two-sided sign-flip permutation tests with TFCE. We quantified the tested effect as a location-by-time map for each observation contributing to the group-level test, such as the within-subject standardized slopes *S*_(_ in Eq. (**2**). In each permutation, we randomly multiplied each entire effect map by (+1) or (-1), preserving the spatial and temporal structure within each map, and recomputed the corresponding group-level TFCE map. We recorded the maximum absolute TFCE score across the electrode–time map for each permutation to control the family-wise error rate. For each observed TFCE, we calculated the family-wise-error-corrected permutation p-value for each location *l* and time *t*:

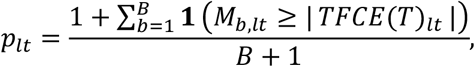

where *B* = 4,000, *M*_*b*_ is the maximum absolute TFCE score in permutation *b*, and *TFCE*(*T*)_*lt*_ is the observed TFCE score.

### Simulation-based power analysis to detect association between visual ERP amplitude and task difficulty

Our cluster-based permutation testing showed no significant association between visual ERP amplitude and working memory load. To determine the magnitude of the association that could be detected with our sample size and statistical analysis, we performed a simulation-based power analysis. Let **X**^*s, j*^ ∈ ℝ^*I*×*T*^ denote the electrode-by-time matrix of visual ERP amplitude, for subject *s* and difficulty level *j*. We first constructed a null dataset 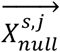 by removing the observed linear association between task difficulty and ERP amplitude separately for each subject:

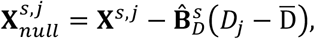

where 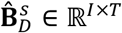 contains the estimated coefficient 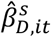 for all electrodes and time windows. To obtain 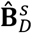, we modeled visual ERP amplitude for electrode *i*, and time *t* as

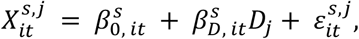

where *D*_*j*_ codes task-difficulty level, 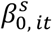 denotes the intercept, 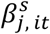 denotes the observed linear difficulty effect, and 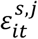 denotes the remaining level-to-level variation.

For each simulation, we generated a bootstrap sample of 20 subjects by sampling 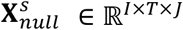 with replacement, preserving the observed within-subject covariance structure. We then injected difficulty-related ERP effects into two electrodes (i.e., C3, C4) from 0.30 to 0.40 s. To preserve between-subject variability, for each simulated subject, we sampled a subject-specific effect :

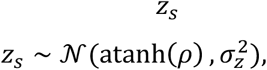

where *ρ* is the assumed Pearson correlation ranging from 0.1 to 0.5; and *σ*_*z*_ is the observed between-subject standard deviation of Fisher-*z*-transformed difficulty–ERP correlations within 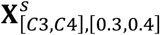. We then obtained the simulated correlation effect by the inverse Fisher transformation *r*_*s*_ = tanh(*z*_*s*_). We next generated simulated visual ERP amplitudes by adding simulated correlation effect 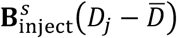 to the empirical null data:

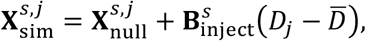

where 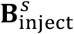 represent the correlation effect at visual ERP amplitude unit

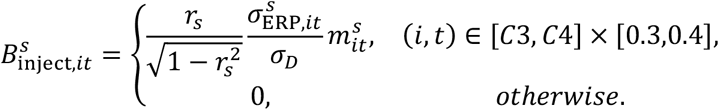

*σ*_*D*_ is the empirical variability of 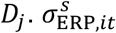 denotes the standard deviation of 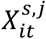 across difficulty levels for subject *s*, electrode *i*, and time 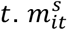 represents the variability of the injected effect magnitude across (*i, t*) ∈ [*C*3, *C*4] × [0.3,0.4] within a subject *s*:

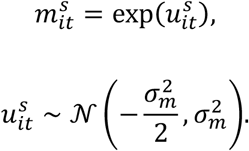

We sampled 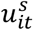 from a Gaussian distribution with mean 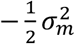, which ensures that 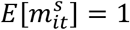. Therefore, 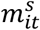 does not systematically increase or decrease the average injected effect. We set *σ*_*m*_ to 0.1 to reflect modest variability.

To estimate power, we repeated cluster-based permutation tests with TFCE on **X**_sim_. Briefly, for each subject *s*, electrode *i*, and time *t*, we regressed the simulated ERP values 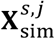 across difficulty levels *D*_*j*_. The estimated slope is

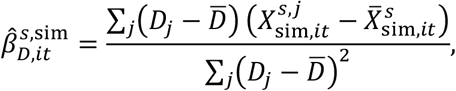

where 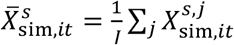.

We performed TFCE on t-like statistics based on 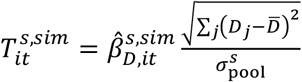. 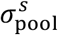 is a subject’s residual visual ERP variability across difficulty levels, electrodes, and time. To calculate 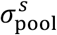 empirically, we normalized visual ERP amplitude using z-score within each electrode. We then fit the difficulty regression separately for each electrode and time and pooled the. We then calculated 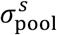 for subject *s* as the median absolute deviation of resulting residuals across all difficulty levels, electrodes, and time points.

For each simulation, a simulated effect was considered detected if ∃(*i, t*) ∈ [*C*3, *C*4] × [0.3,0.4] reached TFCE-corrected *p* < 0.05. For each candidate correlation *ρ*, we estimated the statistical power as 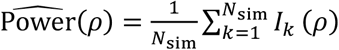, where *N*_sim_ is the number of simulated datasets and *I*_*k*_ (*ρ*) = 1when the injected effect was detected in simulation *k*, and 0 otherwise.

### Source localization

Visual ERPs recorded from scalp EEG reflect the combined contributions of distributed cortical sources, limiting our anatomical interpretation. To generate hypotheses about cortical regions involved in vtaVNS-related modulation of visual processing for subsequent testing with intracranial EEG in patients with epilepsy, we estimate visual responses in cortical source space. For each participant and condition, we pooled visual stimulus-locked epochs across stimuli and difficulty levels, calculated their median, and baseline-corrected resulting waveforms by subtracting their mean amplitude during the 500 ms preceding stimulus onset.

To localize cortical sources, we used standardized low-resolution brain electromagnetic tomography (sLORETA), implemented in Brainstorm^47,70^. sLORETA estimates cortical sources using a standardized minimum-norm solution. We estimated the source signal at 100 Hz for computational efficiency. For healthy participants, we placed EEG electrodes according to the standard 10–10 system following the American Clinical Neurophysiology Society guidelines. To define the forward model. we used Brainstorm’s ICBM152 template head model with standard 10–10 electrode positions. The cortical source space comprised 15,002 vertices, with source orientations constrained perpendicular to the cortical surface. To account for differences in background variability across electrodes and correlations between their signals, the inverse solution incorporated a noise covariance matrix. For each participant and condition, we estimated noise covariance from the pre-stimulus baseline.

To identify candidate anatomical locations where vtaVNS reduced visual responses for subsequent intracranial recording testing, we compared the magnitude of source activity between the vtaVNS and sham conditions across cortical regions and time. We quantified magnitude irrespective of source direction because our question concerned the strength of visual response. Within each participant, condition, and cortical vertex, we standardized source waveforms using their prestimulus mean and standard deviation, averaged the standardized signed waveforms across participants, and calculated their absolute values. Negative contrasts indicate smaller response magnitudes during vtaVNS. To characterize anatomically and temporally consistent patterns, we applied local spatiotemporal consistency weighting. Let *x*_*l,t*_ denote the source-activation contrast at location *l* and time *t*. For each *x*_*l,t*_, we defined a local neighborhood *N*_*l,t*_ containing samples from the same Desikan–Killiany anatomical region within a ±50-ms temporal window. We first estimated the local effect at each point *m*_*r,t*_ as the 10% trimmed mean of neighboring values with the same sign as *x*_*r,t*_. We quantified how closely the magnitude of the observed contrast matched this local effect using a Gaussian similarity function 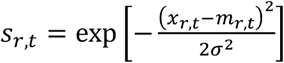, where *σ* was set to 25% of the robust global contrast scale (the 98^th^ percentile of |*x*|). Thus, *s*_*r,t*_ approached 1 when the local contrast had a magnitude similar to that of its surrounding same-sign pattern, and decreased as its magnitude diverged from that pattern. We down-weighted contrasts whose magnitudes differed substantially from their local neighborhood

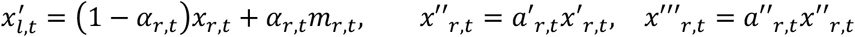

where *α*_*r,t*_ = 0.70*s*_*r,t*_ *a*′_*r,t*_ = 0.25 + 0.75*s*_*r,t*_, *a*′′_*r,t*_ is the proportion of the neighborhood’s absolute effect magnitude having the same sign as *x*_*l,t*_. In *x*′′′_*r,t*_, vtaVNS-sham contrast that closely matched the surrounding pattern was largely preserved.

We next used *x*′′′_*r,t*_ to identify spatiotemporal clusters of consistent vtaVNS-related reductions in the visual response. To achieve this, we identified negative *x*′′′_*r,t*_ within the top 2% of effect magnitude as seeds. We then progressively lowered the threshold and added only samples that remained connected to an existing seed. We defined connectivity across anatomical locations and neighboring time samples within ± 10 ms. We retained clusters spanning at least 50 ms and two anatomical locations. For each retained cluster, we quantified its integrated absolute effect magnitude as 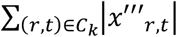, where *C*_*k*_ denotes the cluster.

To visualize cortical sources over a given time interval, we averaged the source activation magnitude at each cortical vertex over the interval (e.g., 200–600 ms after stimulus presentation). We generated a triangulated cortical surface from the Brainstorm head model using the *Matplotlib* Python package. We color-coded each surface triangle based on the mean of source activation magnitude at its three vertices.

### Quantification of distance to criticality

In dynamical systems, criticality describes an operating regime in which neural activity is neither strongly damped nor excessively amplified. This operating regime provides a plausible network configuration for efficient cortical processing. Under a critical operating regime, cortical dynamics are highly tunable, allowing the brain to flexibly reconfigure activity patterns to meet changing computational demands^49^. Our cortical-efficiency hypothesis predicts that vtaVNS shifts cortical dynamics toward this critical operating regime.

In this section, we first describe the physical intuition and mathematical framework that characterize critical dynamics in a time series, and then detail our implementation for quantifying distance to criticality. The mathematical framework for critical autoregressive (AR) processes has been developed previously^71^. Here we also focus on AR processes because they are established in EEG analysis for characterizing behaviorally and physiologically relevant temporal dynamics, while also permitting an analytical solution to quantify distance to criticality^72,73^. Briefly, critical dynamics are scale invariant, meaning that they are not characterized by a single temporal scale. When critical dynamics is progressively coarse-grained, its statistical structure is preserved across scales. In contrast, noncritical dynamics progressively lose their temporal correlation structure and flow toward a trivial fixed point.

To formalize the idea of critical dynamics, we first describe the trajectory probability distribution of an AR process and express it in Fourier coordinates, in which fluctuations at different temporal scales are separated by frequency. We then define the temporal coarse-graining operator, which marginalizes out high-frequency Fourier components while retaining slow dynamics. A defining property of critical dynamics is that the trajectory probability distribution remains unchanged, which we used to quantify how far an observed cortical time series lies from the critical regime.

An order-*p* AR process can be expressed as:

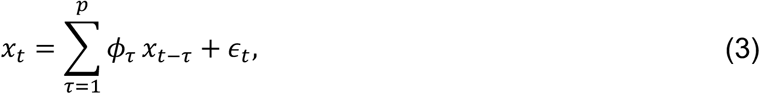

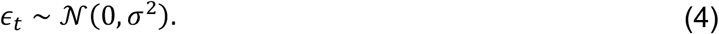

AR model defines a probability distribution *P*[*x*] over entire time-series:

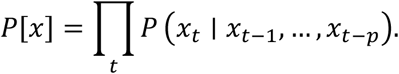

From Eqs. (**3**) and (**4**) and because of the additive of Gaussian distribution, we have

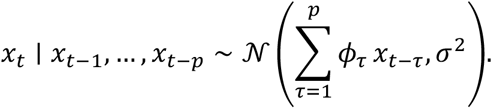

Therefore,

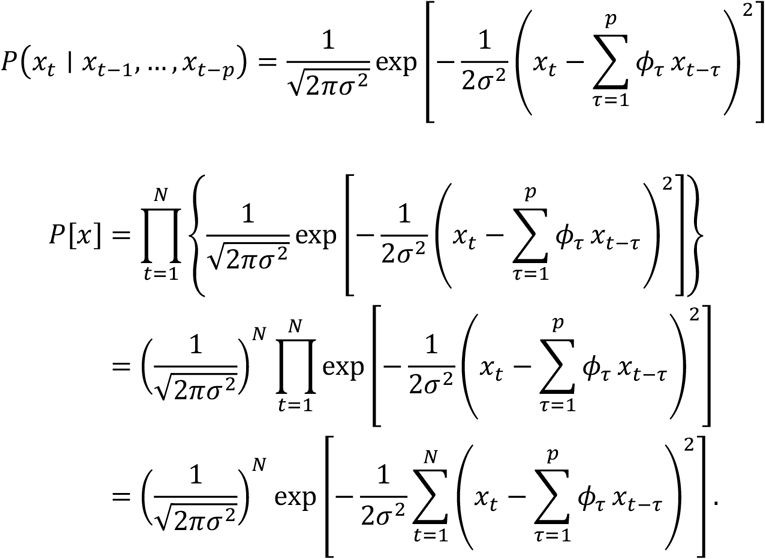

Thus,

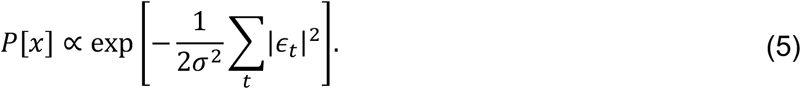

Temporal coarse-graining integrates out the high-frequency dynamics. To formally define the temporal coarse-graining operator, we transformed the AR process into the frequency domain, where fast and slow temporal fluctuations are represented by high- and low-frequency components, respectively.

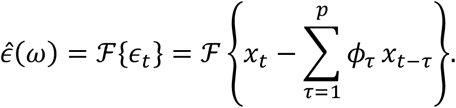

Using the time-shift property 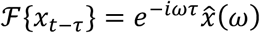, we obtained

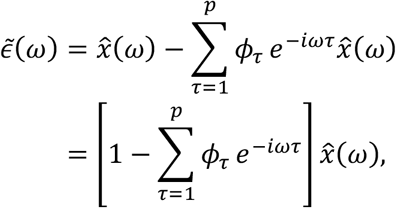

where 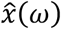 is the Fourier transform of *x*_*t*_. Using Parseval’s theorem, we have

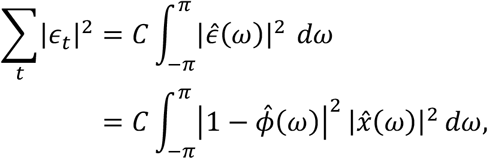

where 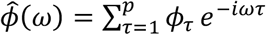.

From Eq. (**5**), we have

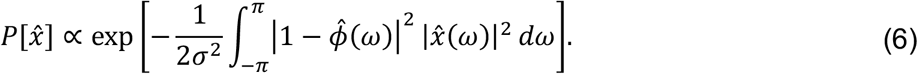

Temporal coarse-graining examines the dynamics at progressively longer timescales. We define the temporal coarse-graining operator *ℛ*: *P*[*x*] → *P*′[x′], which integrates out Fourier components above a cutoff *ω*_*c*_:

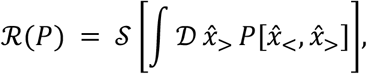

where 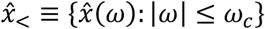 denotes the collection of low-frequency Fourier components of the trajectory retained after coarse-graining, 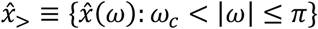, and 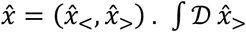 integration over the amplitudes of all high-frequency Fourier modes. As a notation 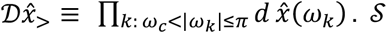 denotes a rescaling operator, allowing us to compare the trajectory probability distribution at a coarser temporal resolution. From Eq. (**6**), we can write

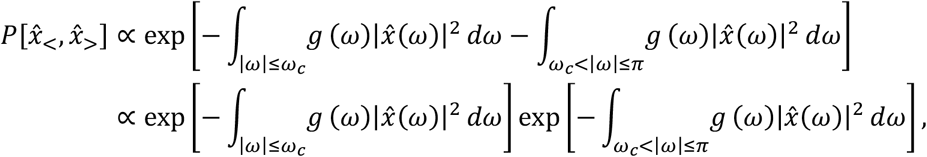

where

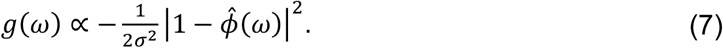

Thus,

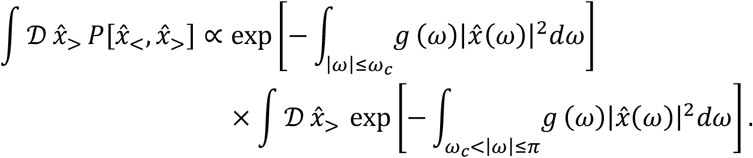

If the Fourier representation has a finite set of discrete frequency coefficients 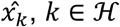, the second term can be expressed as follows. *H* is a finite set of high-frequency Fourier indices.

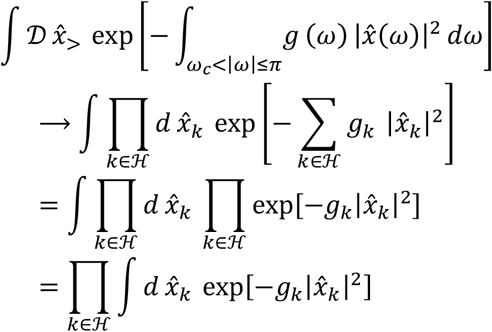

Note that

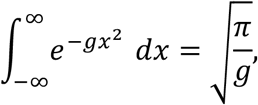

The second term is a constant; thus,

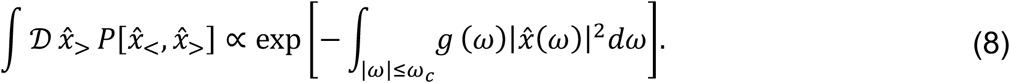

The rescaling operator *S* sets the frequency range back to [−*π, π*], *by setting*

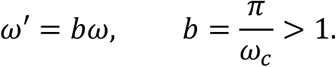

From Eq. (**8**)

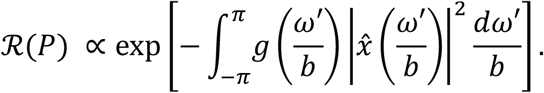

If we set

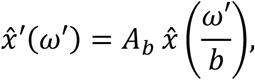

then,

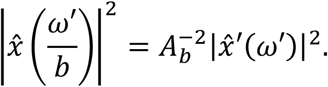

If we set

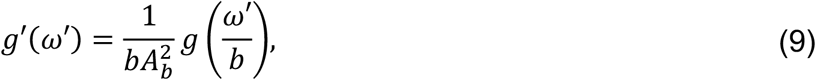

we have 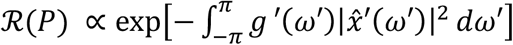|

Critical dynamics correspond to kernels whose functional form is unchanged under coarse-graining, that is, *g*′ = *g*. We will use an AR(1) process to illustrate critical and noncritical dynamics. For *x*_*t*_ = *ϕx*_*t*−1_ + *ϵ*_*t*_, the kernel is 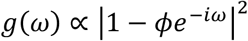. We can expand the expression:

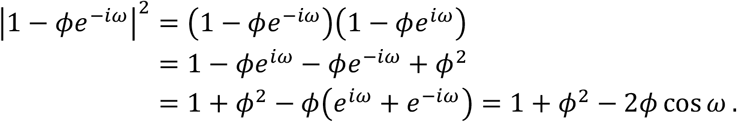

Near *ω* = 0, Taylor expansion of cos *ω* yields *g*(*ω*) = (1 − *ϕ*)^2^ + *ϕω*^2^ + *O*(*ω*^4^). Let *ϕ* = 0.5, *g*(*ω*) ≈ 0.25 + 0.5*ω*^2^. From Eq. (**9**), under repeated coarse-graining and given *b* > 1,

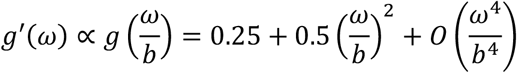

Repeated temporal coarse graining gives 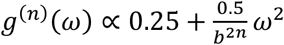. We previously showed

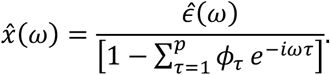

The power spectral density takes the following form:

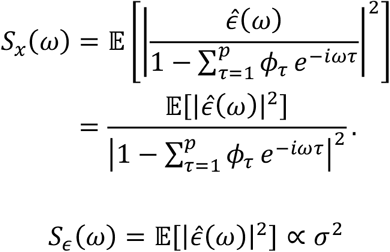

Thus,

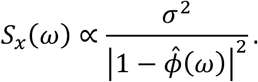

Comparing with Eq. (**7**), we see

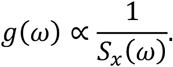

From the Wiener–Khinchin theorem,

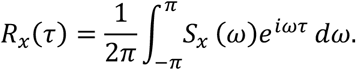

If 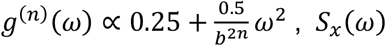 would approach a constant, hence autocorrelation *R*_*x*_ (*τ*) = 0. This suggests that repeated temporal coarse graining yields temporally uncorrelated dynamics (i.e., white noise).

In the case of critical dynamics, let 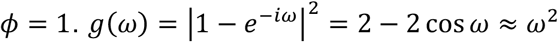. After coarse graining, the functional form *g*(*ω*) is preserved. Thus, the statistical probability is temporally scale-invariant.

To generalize from these two examples, we can expand Eq. (**7**) near *ω* = 0:

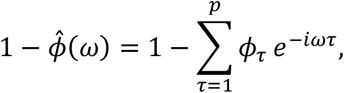

and

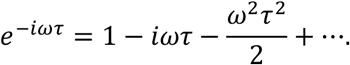

Thus,

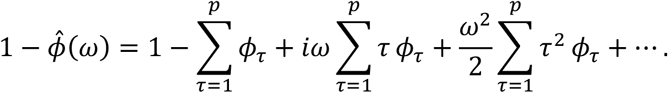

When 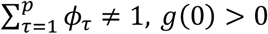. Near *ω* = 0, *g*(*ω*) = *g*_0_ + *O*(*ω*). Under repeated temporal coarse-graining, the AR process would become temporally uncorrelated dynamics, as shown in the first example.

Therefore, the manifold M_*c*_ = {***ϕ*** ∈ ℝ^*p*^: **1**^⊤^***ϕ*** = 1} defines the critical hyperplane for the AR process. We define distance to criticality as 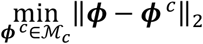. Operationally, we calculated distance to criticality as the shortest Euclidean distance the fitted AR coefficient vector and the convex hull spanning critical AR coefficient vectors *v*^(N)^, following the approach proposed and validated by Lim et al.^38^ We divided each trial into five consecutive, nonoverlapping 0.5-s windows. Each trial contains EEG signal from -1 to 1.5 s relative to stimulus presentation. For each window,

we fitted an AR (25) model to the amplitude envelope of the mean-centered signal *A*(*t*) using the Yule-Walker method. To construct the critical hyperplane manifold M_*c*_, we used AR coefficient vectors corresponding to characteristic polynomials (1 − *L*) ^*m*^, for *m* = 1, …, 25, where *L* denotes the lag operator such that *Lx*_*t*_ = *x*_*t*−1_, *L*^2^*x*_*t*_ = *x*_*t*−8_.

These AR processes lie on the critical manifold M_*c*_ because they can be expressed as

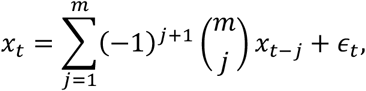

and

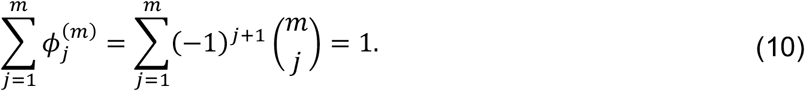

Eq. (**10**) is obvious when we replace L with 1 in 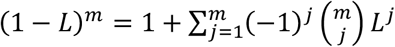; thus 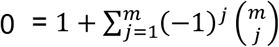. For example, for *m* = 2, the AR process is (1 − 2*L* + *L*^2^)*x*_*t*_ = *ϵ*_*t*_, that is, *x*_*t*_ = 2*x*_*t*−1_ − *x*_*t*−2_ + *ϵ*_*t*_. For a critical AR process, its coefficient vector 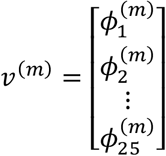. We define the convex hull *Vw* as the set of all weighted averages of the critical vectors, where the weights are nonnegative and sum to 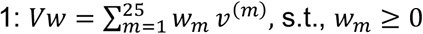 and 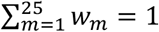.

We therefore calculated the distance to criticality for each 0.5-second window as 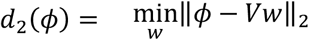. The averaged distance to criticality for the 5 non-overlapping windows is the distance to criticality for each trial.

### Assessing the effect of vtaVNS on distance to criticality

To test a prediction of the cortical-efficiency hypothesis that vtaVNS shifts cortical dynamics toward a more critical operating state, we assessed how distance to criticality varied with memory encoding outcome, task difficulty, and stimulation condition using a multivariate linear model. For each subject, difficulty level, memory encoding outcome (successful versus unsuccessful), and electrode, we averaged distance to criticality across trials. Let **d** denote the vector of distance to criticality containing the distance-to-criticality value for each electrode. We fitted the following linear regression using ordinary least squares:

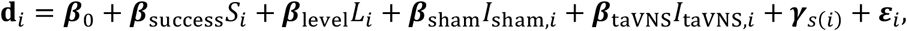

where **d**_*i*_ is the vector of distance-to-criticality values for observation *i, S*_*i*_ codes memory encoding outcome, *L*_*i*_ is difficulty level, *I*_sham,*i*_ and *I*_taVNS,*i*_ indicate sham and vtaVNS conditions, and ***γ***_*s* (*i*)_ accounts for between-subject differences in distance to criticality. For each predictor, the estimated ***β*** contains coefficients for each electrode. We estimated regression coefficients and assessed their statistical significance using a within-subject Freedman–Lane permutation procedure. For each effect, we generated a null distribution that represented the magnitude of the estimated effect expected if the corresponding null hypothesis were true. Specifically, we first fitted a restricted version of the regression model in which the effect under test was constrained to zero; for example, we imposed ***β***_*taVNS*_ − ***β***_*sham*_ = 0 to test the effect of vtaVNS. We retained the fitted values from the restricted model and permuted its residuals across observations within each subject. Because each observation contained residuals from all electrodes, we permuted the entire residual vector, preserving the covariance structure across electrodes. We added the permuted residual vectors back to the restricted-model fitted values to generate a dataset under the null hypothesis and then refitted the original, unrestricted model. For each permuted dataset, we estimated the effect vector corresponding to the hypothesis being tested: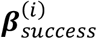 for memory encoding success, 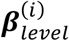 for difficulty level, and 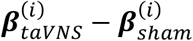 for the vtaVNS-versus-sham. We repeated this procedure 5,000 times, thus *i* = 1,2,3, …, 5000. We summarized the magnitude of each effect vector using its Euclidean norm, for example, *T* = *‖****β***_*taVNS*_ − ***β***_*sham*_*‖*_2_. We calculated the p-value as the proportion of null statistics greater than or equal to the observed statistic.

### Electrode implantation and clinical procedure

All patients underwent implantation of multi-contact depth electrodes (sEEG Depthalon depth electrode, PMT® Corporation, MN, USA) for seizure monitoring. Each electrode had 8–16 contacts (2 mm cylindrical contact length, 0.8 mm diameter, spaced 3.5 mm apart center-to-center).

The clinical team planned the electrode placement to ensure appropriate coverage of the brain for seizure network localization. T1-weighted imaging (MPRAGE sequence, voxel size

1.0 × 1.0 × 1.0 mm^3^) was obtained in the presurgical workup and loaded into the planning workstation. The neurosurgeon implanted electrodes, guided by a stereotactic robotic system (ROSA ONE Brain, ZIMMER BIOMET, IN, USA). After surgery, patients underwent a CT scan, which was co-registered to their pre-operative MRI to obtain precise 3D coordinates for each electrode contact. We describe the methods for electrode localization in **Method: Intracranial electrode localization**. Patients were then awakened from general anesthesia and transferred to the Epilepsy Monitoring Unit (EMU) for intracranial electroencephalography (iEEG) monitoring. During their EMU stay, the research team introduced the study to the patients and obtained written informed consent before beginning any experimental recordings. Following consent, research sessions were scheduled in coordination with the clinical team to ensure no disruption to standard clinical care. Standard clinical protocols may taper or withdraw antiseizure medications to provoke seizures during iEEG monitoring. In some cases, partial sleep deprivation was employed to increase seizure likelihood.

All electrophysiological data were collected using the EMU’s clinical intracranial neural recording system and other devices that are FDA-approved for sale for studies in human subjects or are commonly approved and used at other research institutions.

### Intracranial electrode localization

We related intracranial electrophysiological recordings to participant-specific anatomy using an automated pipeline that integrated postoperative computed tomography (CT), preoperative structural magnetic resonance imaging (MRI), anatomical segmentations, and electrode trajectories exported from the ROSA robotic surgical system. We were motivated to develop an automatic pipeline because manual electrode localization is time-consuming and can introduce operator-dependent variability. Our central strategy was to combine the strong CT contrast produced by metallic electrode contacts with prior knowledge of each electrode array’s geometry and contact configuration to obtain reproducible estimates of contact locations. Our pipeline consisted of three main steps. First, we transformed all imaging and surgical electrode-trajectory information into a common participant-specific anatomical space defined by the preoperative MRI. Second, we identified individual electrode arrays on the postoperative CT and estimated their contact locations from the CT signal and the known geometry of each array. Third, we linked the localized contacts to electrophysiological recording channels using participant-specific channel-mapping tables generated during electrode connection in the operating room and assigned anatomical labels from the MRI-derived brain segmentation.

### Image coregistration

The ROSA surgical assistive system contained the spatial coordinates and trajectories of the implanted electrode arrays. To coregister postoperative CT and the MRI volume exported from the ROSA surgical planning system, we converted all volumetric images to right–anterior–superior (RAS) orientation and converted the MRI and CT scans from DICOM to NIfTI format using dicm2nii from the NeuroImaging Tools and Resources Collaboratory. To ensure robust coregistration using SPM12, we rigidly translated the CT volume so that its geometric center was located at the world-coordinate origin. We rigidly coregistered volumetric images to the preoperative MRI using the mutual-information-based registration implemented in SPM12.

### Identification and isolation of individual electrode arrays

The goal of identifying and isolating individual electrode arrays is to estimate the position of each individual contact. Our approach is to first construct a spatial model of each implanted electrode array and then combine this model with the known array specifications, including contact length, contact spacing, and number of contacts, to estimate contact positions along the array. Constructing the electrode array model required identifying the CT voxels belonging to each electrode array and fitting a geometric trajectory to them. In this section, we describe methods to identify such CT voxels and to separate them from neighboring electrode arrays.

Because metal is highly attenuating on CT, electrode contacts and other metallic structures appear as high-intensity voxels. We first identified CT voxels likely to correspond to the metallic components of the implanted electrode array. To do this, we estimated the positive-intensity distribution and identified a valley separating the two most prominent high-intensity peaks, yielding a threshold beyond which we consider metallic voxels. The remaining challenge was to determine which metal voxels belonged to each individual electrode array. For each implanted array, we used the executed electrode trajectory recorded in the ROSA system to define a spatial search region comprising voxels within 4 mm of the trajectory. Within a search region, we used a multi-seed-growing procedure to identify a spatially coherent set of metal voxels corresponding to the target electrode array. We selected 6 seeds from local CT-intensity maxima after Gaussian smoothing with a standard deviation of 0.6 voxels. We ranked seeds using a weighted combination of normalized CT intensity and proximity to the ROSA trajectory, and required them to be separated by at least 5 mm. Each seed independently initialized a growing set of voxels containing up to 8 nearby candidate voxels.

Starting from each seed, we iteratively added candidate metal voxels to construct a voxel set representing a single electrode array. Each seed initialized an independent voxel set containing up to eight nearby candidate voxels. At each iteration, we evaluated a candidate voxel according to two principles: spatially connected to the existing set, and its addition preserve the approximately linear, elongated geometry expected for an implanted depth electrode.

To quantify this geometric consistency, we enclosed the current voxel set within a cylindrical tube aligned with the candidate electrode axis and calculated a filling fraction, defined as the total volume of selected voxels divided by the volume of the enclosing tube. We adjusted the tube radius to span the spatial extent of the selected voxels, with the axis permitted to vary by up to 3 degrees from the ROSA-derived trajectory direction. Voxels belonging to the same electrode array tended to increase the occupied volume while requiring little expansion of the enclosing tube. In contrast, adding a voxel from a neighboring electrode or other metallic structure typically increased the tube volume disproportionately, thereby reducing the filling fraction. These principles defined our voxel-inclusion rules. We rejected a candidate voxel if it was more than 3 mm from the existing voxel set, if its addition decreased the filling fraction by more than 0.2, or if the resulting filling fraction fell below 0.05. Growth continued until no remaining candidate voxels were left. Because region growing was initialized independently from six seeds, the procedure produced six candidate electrode voxel sets for the same array. We combined these results by retaining voxels that appeared in at least two independently grown sets, yielding a consensus electrode mask.

The computational cost and memory requirement grow with the resolution of CT images. We performed the seed-growing procedure on subsampled CT images when the CT slice thickness was *≤* 0.5 mm. We then mapped the consensus electrode mask back to the original CT grid. To further reduce contamination from neighboring metallic structures, we applied a global directional-coherence measure. For each retained voxel, we assessed whether its spatial relationship to the remainder of the mask was consistent with a common electrode axis. We constructed unit vectors from each voxel coordinate to the 50% farthest voxels and calculated directional coherence as the mean absolute cosine similarity among all unique pairs of these vectors. Values closer to 1 indicated that the vectors were predominantly aligned along a common axis, as expected for voxels belonging to the same electrode array. We used the absolute cosine similarity; thus, vectors pointing in opposite directions along the same electrode axis were considered equivalent. We removed voxels with coherence values below 0.98 from the electrode mask.

### Modeling electrode array

After extracting the CT voxels associated with each electrode array, we converted this voxel set into a geometric model of the implanted array, which was subsequently combined with the known electrode specifications, including contact length, intercontact spacing, and number of contacts, to constrain estimation of individual contact locations. In most cases, the implanted depth electrodes are approximately linear but can be bent in rare circumstances. We considered both a straight-line model and a quadratic Bézier-curve model and selected the representation that better described the observed CT voxels without unnecessary model complexity.

A straight-line model is parameterized by direction and two endpoints. To estimate the direction, we used a FreeSurfer-derived brain mask to distinguish metal voxels within the brain from extracerebral metal and then fitted a straight centerline to the set of intracranial voxels associated with individual electrode arrays. We defined the initial axis as the principal direction of the voxel coordinates using Principal Component Analysis and refined it by iteratively reweighted fitting of perpendicular point-to-axis distances using Tukey biweight weights. To estimate the endpoints for an electrode array, we decomposed voxel coordinates relative to the fitted centerline into a radial distance and an axial position along the axis. We estimated the endpoints based on axial coordinates of voxels within the 90^th^ percentile of the radial-distance distribution. we used the 98.5th percentile of the axial distribution at each end to obtain robust endpoint estimates. For the proximal endpoint, we constrained the estimate to remain within the plausible extracerebral portion of the implanted lead. We required the proximal endpoint to lie on the proximal side of the brain-entry point and within a radial distance of 2 mm.

To accommodate potential electrode curvature, we also fitted a quadratic Bézier curve with fixed proximal and distal endpoints and a single free three-dimensional control point. We optimized the control point by minimizing squared point-to-curve distances after excluding the lowest and highest 5% of residuals, to reduce sensitivity to potential metal artifacts. We compared the straight and curved model using a Bayesian information criterion(BIC)-inspired criterion:

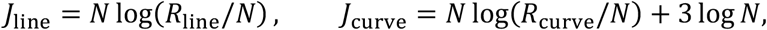

where *N* is the number of intracranial points and *R* is the sum of point-to-model distances after trimming 5% from each tail. 3 log *N* is the BIC-like complexity penalty for the three additional parameters needed in the Bézier curve model. All electrode arrays were better modeled linearly.

### Geometry-constrained electrode contact localization

We localized individual electrode contacts by fitting the known contact configuration along the geometry model of electrode arrays to best explain the observed CT signal. The geometric model spatially constrained candidate contact locations. The localization problem was reduced from unconstrained three-dimensional estimation to a one-dimensional optimization along the modeled trajectory. We obtained the contact configuration from operative metadata and cross-validated it against data in the ROSA surgical assistive system. Contact configuration includes the electrode model, serial number, and contact count. These specifications determined the expected contact dimensions and relative spacing along the array. For example, PMT depth electrodes had a nominal diameter of 0.8 mm and contact length of 2 mm, with center-to-center offsets determined by the corresponding electrode model. We represented the complete contact arrangement as a rigid geometric template and optimized its position along the electrode trajectory. Fitting all contacts jointly preserved their known relative spacing and allowed CT evidence from the full array to contribute to localization, improving robustness when individual contact boundaries were blurred or poorly resolved.

To determine where along the centerline the contact template best aligned with the observed CT, we defined a contact-evidence score *p*(*s*) at successive positions *s* along the trajectory. The score was based on the physical expectation that a metallic contact should produce higher CT intensity than the adjacent insulating portions of the lead and surrounding tissue. At each candidate position, the contact-evidence score compared CT intensity within a cylindrical core at this location with the intensity in a surrounding shell. For each electrode array, we defined the core radius as the larger of 0.5 mm and the 97.5th percentile of radial distances between voxels and the centerline of the electrode array model, after excluding voxels within the proximal and distal 5% of their axial distribution. The core extended axially to ±0.6 contact lengths around the candidate position.

The surrounding shell included a radial annulus around the candidate contact and two full-disk regions extending axially to the insulating intervals between neighboring contacts. Let *s*_0_ denote position along the electrode array centerline, and *u* denote axial position relative to the center of the candidate contact. We defined the associated contact-evidence score as

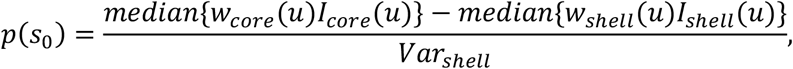

where *I*_*c*_ and *I*_*s*_ denote CT intensities from the core and shell, respectively, and *w*_*c*_ and *w*_*s*_ are axial weights assigned according to each sample’s position relative to the candidate contact center. *Var*_*shell*_ is a robust estimate of local intensity variability in the surrounding shell. We applied axial weights to emphasize the portions of the sampling regions that were most informative for distinguishing a metallic contact from adjacent non-contact regions. *w*_*core*_ (*s*) assigned lower weight near the axial boundaries of the core and greater weight toward its central portion:

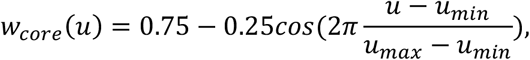

where *u*_*min*_= −0.6, *u*_*max*_ = 0.6.

*w*_*shell*_(*s*) assigned a baseline weight of 0.2 for the annulus surrounding the contact. *w*_*shell*_ (*s*) increased within the proximal and distal insulating intervals:

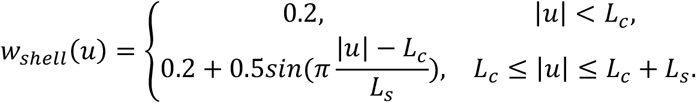

where *L*_*c*_ defines the axial extent of the cylindrical core and *L*_*s*_ defines the length of the adjacent insulating interval sampled on each side.

We subtracted a 20-mm moving-median baseline from *p*(*s*_0_) to reduce slowly varying intensity differences along the lead. We then used the known electrode configuration to jointly estimate the locations of all contacts. We translated the complete manufacturer-specified contact arrangement along the centerline using a common offset. This preserved the expected relative spacing between contacts.

If *L* denotes the total centerline length, *o*_*k*_ the model-specified offset of contact *k* relative to the distal-most contact, and ℓ the contact length, the candidate center position of contact *k* for a common shift *δ* was 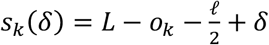. We estimated the common shift by maximizing the summed detrended contact-evidence score across all contacts:

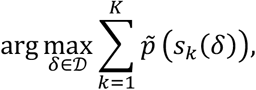

where 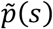 denotes the detrended contact-evidence score, *K* is the number of contacts, and *D* is the set of candidate shifts. This optimization identified the contact configuration that provided the strongest overall agreement with the CT evidence while preserving the manufacturer-specified relative spacing between contacts. We refined the location for each contact within ±0.175 mm, approximately the size of a CT voxel, along the centerline.

### Functional-anatomical location of electrode contacts

After estimating the spatial coordinates of individual electrode contacts, we established the correspondence between each localized contact, its recorded electrophysiological signal, and its participant-specific anatomical location. We then transformed contact locations into a common anatomical space for group-level analysis and visualization.

To link each localized electrode contact to its corresponding electrophysiological recording channel, we used a participant-specific channel-mapping table generated during electrode connection in the operating room. This table recorded the correspondence between the lead identifiers in the ROSA system and the amplifier channel labels assigned by the EEG technician

To determine the brain regions associated with each electrode contact, we performed anatomical brain segmentation using FreeSurfer (v7.4.1), which also creates a cortical and subcortical functional parcellation based on the Desikan-Killiany atlas^74^. We assigned functional-anatomical labels to each electrode contact using a majority-vote volumetric method, in which each electrode was assigned to the brain structure that encompassed the majority of segmented voxels within a 10 mm Euclidean radius of the contact center. We chose a 10 mm Euclidean radius to approximate the spatial reach of local field potential recordings. To assess the consistency of the functional–anatomical labeling of electrode contacts, we performed two complementary analyses. First, we used a surface-based inclusion method, in which electrodes were labeled based on whether their coordinates lay within the triangulated surface mesh of a segmented structure. Second, we implemented a probabilistic labeling approach, in which each structure was assigned a probability proportional to the number of its voxels within a 10 mm sphere centered on the electrode.

For group-level analysis and visualization, we mapped individual electrode locations to a common anatomical space defined by the native MRI of a reference subject. To achieve this, we used the ANTs (Advanced Normalization Tools) image registration framework to coregister each subject’s preoperative T1-weighted MRI to the reference MRI. We applied only the affine component of the transformation to the electrode coordinates to preserve their linear relationships. The affine matrices were estimated within the ANTs’ non-linear optimization pipeline to account for anatomical variability.

### Intracranial EEG data processing

Source localization of the scalp EEG suggested that vtaVNS reduced visually evoked cortical responses in temporal and parietal regions. To test this hypothesis at the level of local neuronal populations, we analyzed intracranial recordings, focusing on broadband high-gamma activity, a marker of population-level cortical activation^42,75^. To remove line noise and volume-conducted activity shared across contacts, we locally re-referenced each contact by subtracting the mean signal of available neighboring contacts within 7 mm. We extracted local high-gamma activity using a fourth-order Butterworth bandpass filter spanning 70–150 Hz, applied forward and backward, and calculated power by squaring the filtered signal. We segmented the high-gamma power into epochs from −1.5 to 1.5 s relative to the onset of each visual stimulus during memory encoding.

To ensure the extracted high-gamma power reflects neuronal responses to visual stimuli, we identified epochs contaminated by movement artifacts or interictal epileptiform discharges, both of which are characterized by abrupt changes in high-gamma power. For each epoch, we z-scored power to its pre-stimulus baseline (-0.5 to 0 s), averaged the z-scored power in 50-ms windows advanced in 25-ms steps, and took the largest absolute difference between consecutive windows as the abruptness score. We excluded epochs with an abruptness score more than six standard deviations above the median score across all epochs, as estimated using the median absolute deviation. We assigned the included epochs to three conditions: vtaVNS (stimulated trials in active vtaVNS sessions), sham (non-stimulated trials in active vtaVNS sessions), and baseline (non-stimulated trials in either active vtaVNS or sham sessions).

### Assessing the effect of vtaVNS on high gamma response following visual stimuli

Source localization identified spatiotemporal clusters of reduced visual response in temporal and parietal regions centered at 0.4-0.5 s. We therefore compared high-gamma amplitude in this window between vtaVNS and sham using linear mixed-effects models of contact-level high-gamma responses. To compute these responses, we subtracted the pre-stimulus mean from each epoch and averaged epochs within each contact and condition. To normalize high-gamma power across contacts without biasing comparisons between conditions, we divided high-gamma power for an electrode contact by the standard deviation of the pre-stimulus samples, pooled across all available conditions for that contact. We fitted separate models for temporal and parietal cortex:

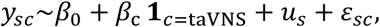

where *y*_*sc*_ is the averaged high-gamma power over the 0.4-0.5 s interval, *u*_*s*_ is a participant-specific random intercept, and *β*_c_ represents the effect of vtaVNS compared to sham.

### Assessing association between successful encoding and reduction in visual ERP

To test whether a smaller cortical response to visual stimuli in a working memory task is associated with successful memory encoding, we compared visual ERP amplitude between successfully and unsuccessfully encoded trials. In the N-back task, participants press a button when the current item matches the item presented *N* trials earlier. The encoding outcome of trial *t* can therefore be read from the response on trial *t* + *N*. We designed our N-back task so that half of the trials are match trials that required a button press. This makes chance-level accuracy 50%: a participant who did not perform the task, or who always (or never) pressed the button, would score 50%.

For each difficulty level, participant and encoding condition, we calculate prestimulus-baseline-standardized visual ERP amplitude at each electrode location and time point. For each task load, we quantified the difference in standardized visual ERP amplitude between the successful and unsuccessful encoding conditions using paired Cohen’s d. We estimated 95% confidence intervals for this effect size (i.e., Cohen’s d) by resampling participants with replacement 10,000 times. The difference in visual ERP amplitude between successful and unsuccessful values for each participant stayed paired during resampling. We assessed significance with a two-sided sign-flip permutation test.

### Quantification and statistical analysis

All data were analyzed and plotted with MATLAB and Python. For regression analyses, we assessed the statistical significance of regression coefficients using two-sided t-tests with degrees of freedom estimated by the residual method implemented in MATLAB’s *fitlme* function. For assessment of statistical significance based on permutation, such as the two-sided sign-flip permutation test used in testing the association between visual ERP amplitude and memory encoding outcome, we computed the p-value as 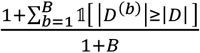, where *D* is the observed statistics (e.g., paired Cohen’s d), 1 is the indicator function, B is the number of permutations, *D*^(*b*)^ is the statistics in *b* ^th^ permutation (e.g., sign-flipping). Statistical significance derived from clustered-based permutation test with TFCE is described in Method: **Cluster-based permutation tests with threshold-free cluster enhancement (TFCE)**. Statistical significance is defined as NS (non-significant), ∗ p < 0.05, ∗∗ p < 0.01, ∗∗∗p < 0.001.

