## Supplementary Materials for "Vibrotactile transcutaneous auricular vagus nerve stimulation (taVNS) during working memory: convergent electrophysiological evidence for cortical efficiency"

### Vibrotactile transcutaneous auricular vagus nerve stimulation promotes efficient cortical processing during working memory: an electrophysiological study

Gansheng Tan *et al.*

*Eric Leuthardt

**This PDF file includes:**

Supplementary Text
Figs. S1 to S4
Tables S1
Movies S1
Audio NA
Data NA
References (1 to 2)

**Other Supplementary Materials for this manuscript include the following:**

Movies S1

### Supplementary Text

#### Validation of artifactual component separation

To validate our approach for separating artifactual components from EEG signals, we examined the fitted template parameters, including scaling and temporal shift, indicating stable artifact morphology (Fig. S1C–D). The median of the scaling parameter is 1.186 for blink component (95% CI = [1.181, 1.191], N = 18317); 0.851 for stimulation artifact component (95% CI = [0.842, 0.862], N = 13539); and 1.263 for cardiac component (95% CI = [1.258, 1.269], N = 51510). The median of temporal shift is 0 for the blink component (95% CI = [-0.833, 0.000]); 0.856 for the stimulation artifact component (95% CI = [0.842, 0.862]); -0.833 for the cardiac component (95% CI = [-0.833, -0.833]). Together, the template-matching procedure separated the neural signal from the blink, stimulation, and cardiac artifact components, providing the basis for the subsequent analyses of visual ERPs.

#### Distance to criticality increases with working-memory difficulty.

To test the cortical-efficiency hypothesis that vtaVNS shifts cortical dynamics toward criticality, we modeled distance to criticality as a function of stimulation condition (vtaVNS versus ssham) while accounting for factors that could influence distance to criticality. Because previous human fMRI and EEG studies have shown that increasing task difficulty suppresses scale-invariant neural activity[1,2], consistent with a shift away from criticality, we used a multivariate linear model with difficulty level, encoding success, and stimulation condition as predictors. Across electrodes, greater difficulty was associated with greater distance to criticality (norm of the electrode-wise coefficient vector = 0.021, permutation p=0.0002; electrode-wise difficulty coefficients >0, range = 0.0019–0.0085 per difficulty level). This pattern is consistent with previous observations that increasing cognitive demand shifts neural dynamics away from a flexible, scale-invariant regime.

#### Behavioral performance during the spatial working-memory task

Our study aims to understand the mechanism by which vtaVNS improves working memory. The primary goal of our spatial working-memory experiment in the patient population was to test the anatomical hypothesis generated from source localization. We analyzed behavioral performance during the spatial working-memory task to verify that patients were able to perform it.

Across participants, the mean accuracy was 78.7% (SD 16.0%), indicating that patients were able to perform the spatial working memory task as instructed. To further explore whether performance differed between active vtaVNS and sham, we categorized trials into 2 intervention conditions: active vtaVNS and sham. Within each intervention condition, we further categorized trials according to the trial-level stimulation state: ON and OFF. We modeled accuracy using a generalized linear mixed-effects model with a binomial distribution and logit link. Trial correctness was the dependent variable, and intervention condition, trial-level stimulation state, and their interaction were fixed effects. A participant-specific random intercept accounted for repeated trials within participants. Active vtaVNS were associated with higher odds of a correct response than sham ($\beta$ = 0.562, Odd Ratio (OR) = 1.75, 95% CI of OR = [1.10, 2.81], p=0.019). Neither the trial-level stimulation-state effect nor the interaction between intervention condition and stimulation state reached statistical significance ( $\beta_{stimulation\ state}$ = 0.430, $p_{stimulation\ state}$ = 0.074; $\beta_{interaction}$ = -0.640, $p_{stimulation\ state}$ = 0.056). This result is consistent with the possibility that the effects of vtaVNS may persist beyond periods of concurrent stimulation. Overall, the behavioral results indicate that patients successfully engaged with the task and actively encoded visual stimuli.

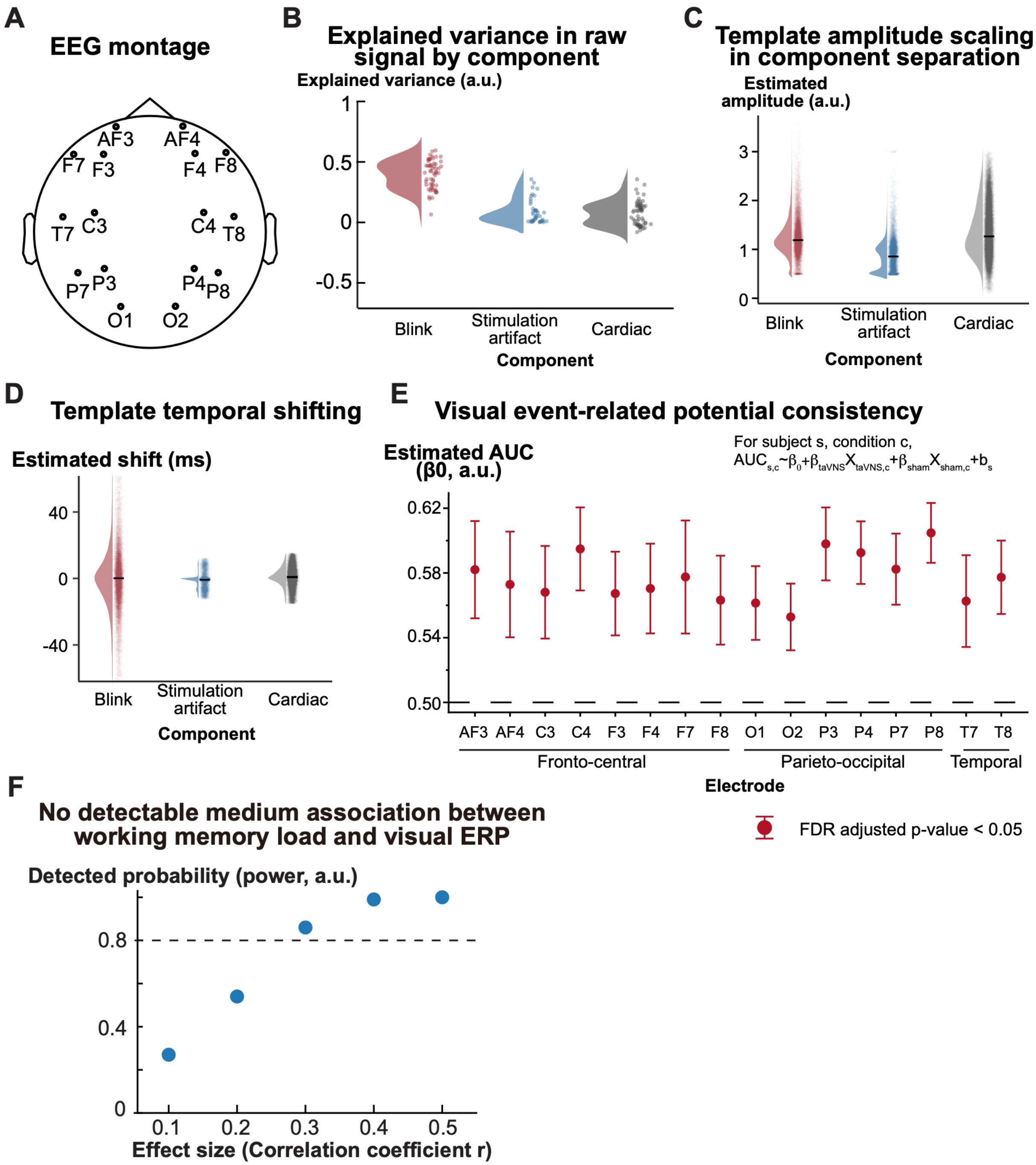


**Fig. S1 (related to Figure 1). Quality control of iterative component separation and identification of visual ERP**. (**A**) EEG montage used in the study. (**B**) Proportion of EEG signal variance explained by the identified blink, stimulation-artifact, and cardiac components. (**C**) Amplitude-scaling coefficient of template fitting to EEG signals. Values centered near 1 indicate that the fitted response has an amplitude similar to the template. (**D**) Temporal-shift coefficient of template fitting to EEG signals. Values centered near 0 indicate no temporal displacement relative to the template. (**E**) Waveform consistency across electrodes, quantified as the area under the receiver operating characteristic (ROC) curve. We constructed the ROC curve by varying the intertrial-correlation threshold and calculating the corresponding true-positive and false-positive rates for distinguishing the post-stimulus interval from the pre-stimulus baseline. Higher area under the ROC curve indicates greater consistency of the visually evoked waveform relative to baseline. This panel shows the mean and standard deviation of the intercept estimates from linear mixed-effects models with area under the ROC curve as the dependent variable and stimulation condition as predictors. (**F**) Post-hoc simulation-based power analysis for detecting a correlation between visual ERP amplitudes and task difficulties. For each simulated dataset, subject-level difficulty effects were recalculated and tested using the same TFCE permutation procedure as the primary analysis (see **Methods**: Simulation-based power analysis to detect association between visual ERP amplitude and task difficulty). We estimated power as the proportion of simulations in which at least one point within the injected electrode–time region reached TFCE-corrected $p < 0.05$.

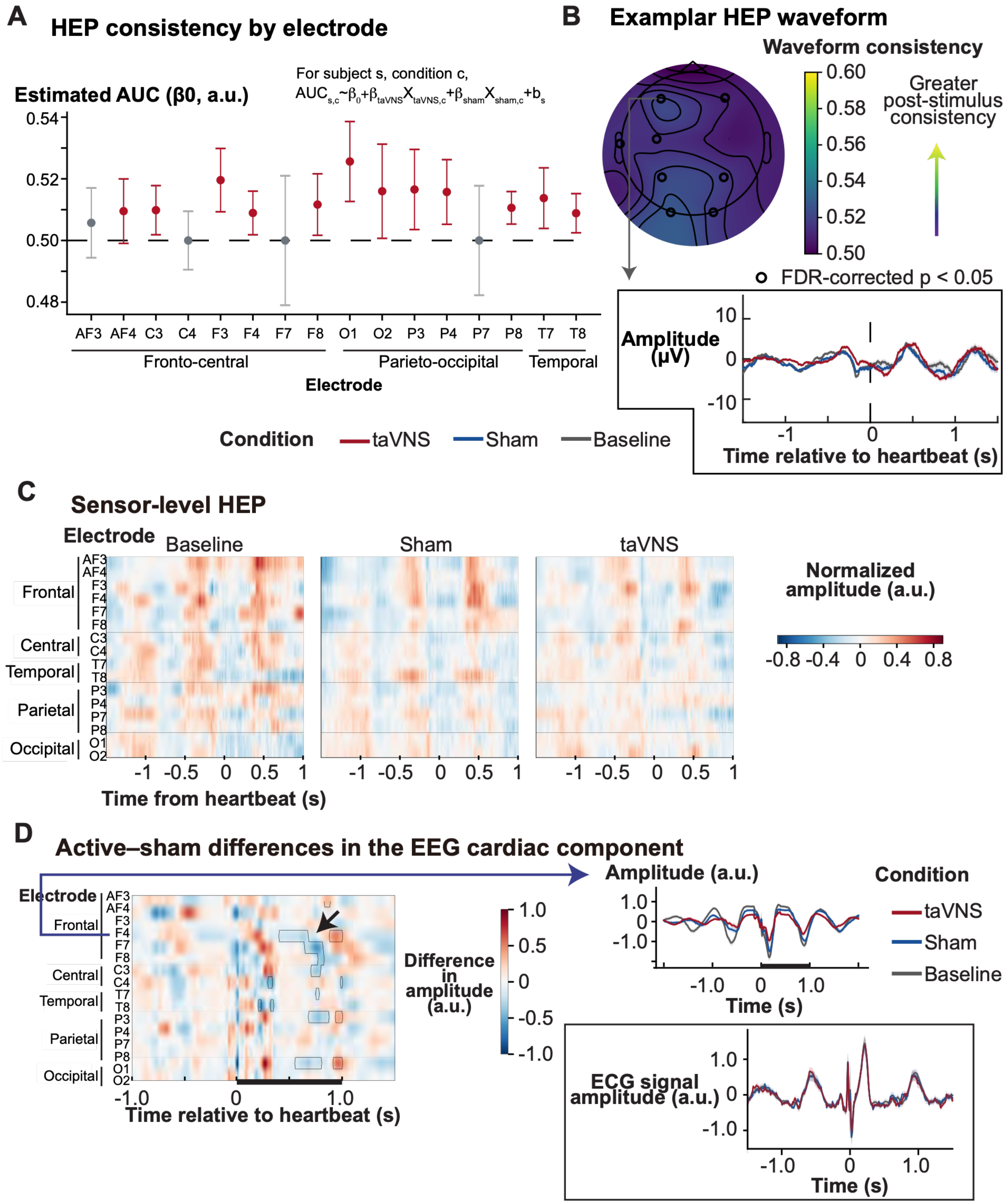


**Fig. S2 (Related to Discussion).** Effect of taVNS on heart-evoked potentials (HEPs). (**A**) Waveform consistency for HEPs extracted from neural components. The error bars represent standard deviation across subjects. (**B**) Cortical distribution of waveform consistency. The inset shows the HEPs for an exemplar electrode during active taVNS, sham, and baseline sections. (**C**)

HEPs across time and electrode during active taVNS, sham, and baseline sessions. Cluster-based permutation test did not identify significant electrode–time clusters in which HEP amplitude differed between active taVNS and sham conditions. (**D**) Difference in cardiac component identified from EEG between active taVNS and sham conditions. The right plot shows the cardiac component signals over time for active taVNS, sham, and baseline conditions. The inset shows the ECG signals for the three conditions.

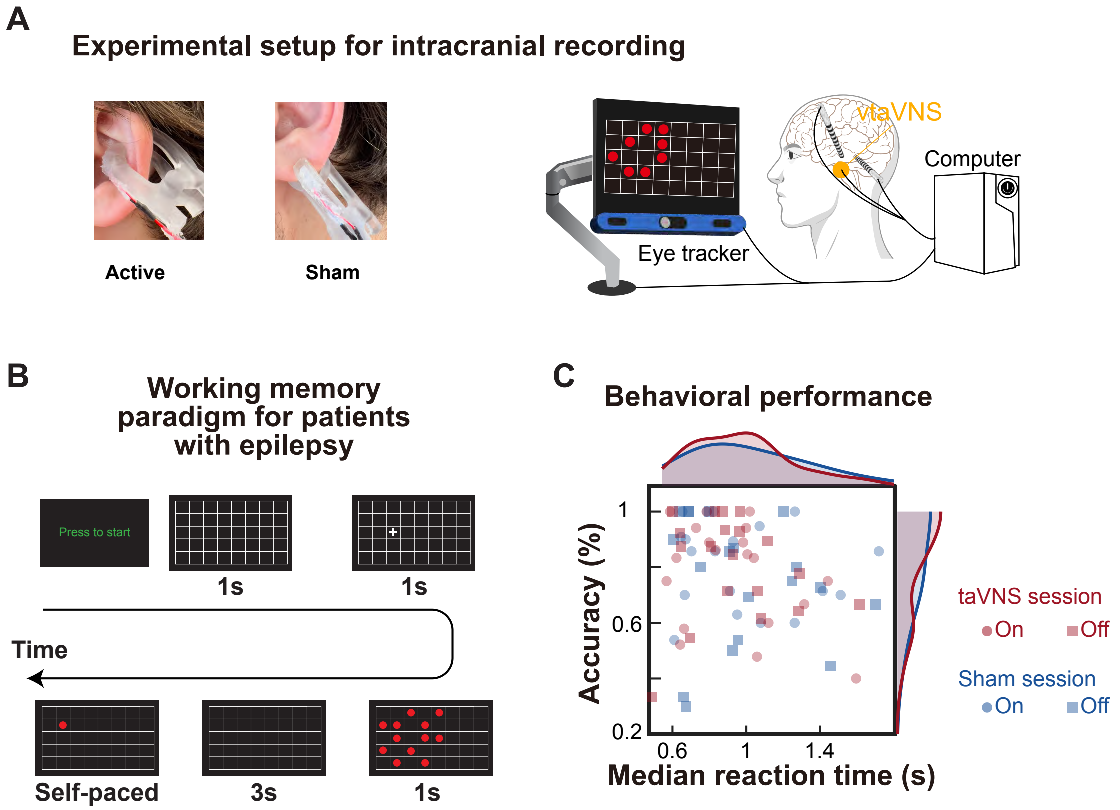


**Fig. S3 (Related to Figure 2)**. Intracranial study of the effect of taVNS on cortical dynamics. (**A**) Miniaturized vibrotactile taVNS device and experimental setup. We used a customized 3D-printed ear clip to house an eccentric rotating mass (ERM) motor and secured the device to the external ear. Active vibrotactile stimulation targets the left cymba conchae, whereas sham stimulation targets the left earlobe. Patients with epilepsy undergoing invasive monitoring in the epilepsy monitoring unit (EMU) completed a spatial working memory task while intracranial local field potentials were recorded. A desktop controls stimulation timing via a digital stimulation box (g.STIMBOX, g.tec medical engineering, Austria), stimulus presentation, and data recording. (**B**) Experimental paradigm. After practice, participants completed multiple sessions of the spatial working memory task. The number of sessions was determined by the clinical schedule and participant availability. Each session consisted of 10 trials, which were randomized to receive either vibrotactile stimulation or no stimulation. Each trial began with the presentation of an empty grid, followed by the appearance of a red cross, which cues participants to the center of the dot pattern during the subsequent encoding phase. During the retrieval phase, participants used a keyboard to indicate whether the dot's location matched that of any dot shown during encoding. To minimize fatigue and vibrotactile taVNS habituation, trials started after the subject pressed a key, with a mandatory 6-second inter-trial interval. (**C**) Behavioral performance during the spatial working-memory task. Each data point represents the accuracy and median reaction time for each subject during taVNS and sham sessions, shown in red and blue, respectively. Marker shape indicates whether stimulation was OFF or ON during the trial within the corresponding taVNS or sham session. Kernel density estimates at the top and right show the distributions of reaction time and accuracy, respectively, color-coded by taVNS and sham session.

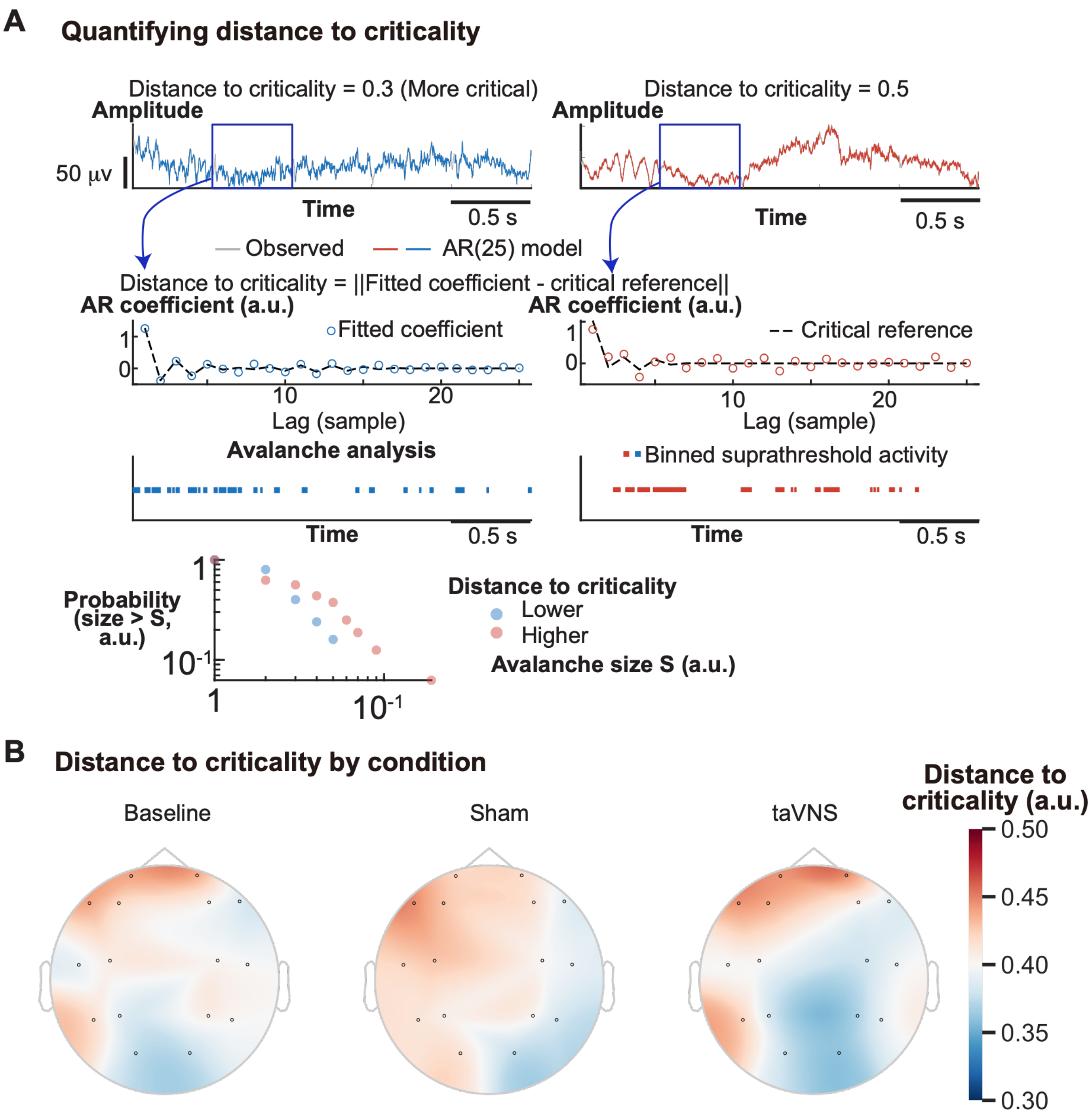


**Fig. S4 (Related to Figure 3).** Quantification of distance to criticality. (**A**) Representative EEG signals with low (left) and high (right) distance to criticality and avalanche analysis of their scale-invariant dynamics. The two plots in the first row show EEG signals (gray) from the predictions from AR(25) models (colored). The two plots in the second row show the fitted AR(25) coefficients and their Euclidean-distance projection onto the manifold of critical AR models for a representative 0.5-s analysis window. The two plots in the third row show binarized temporal bins in which the signal exceeded the burst-detection threshold, defined as the median amplitude plus one robust estimate of baseline variability (i.e., 1.4826 × median absolute deviation). The bottom plot shows empirical cumulative distribution functions of avalanche size (S) for the low- and high-distance-to-criticality examples. Avalanche size refers to the number of consecutive active temporal bins. (**B**) Distance to criticality during baseline, sham, and active taVNS sessions.

**Table S1 Patient Characteristics**

| Patient ID | Age (years) | Sex | Hande-dness | Race | Monitoring duration (days) | Epilepsy diagnosis |
|---|---|---|---|---|---|---|
| 1 | 24 | F | R | Black or African American and White | 10 | Right anterior and posterior hippocampal/basal anterior temporal onset |
| 2 | 22 | M | Ambidextrous | White | 14 | Right frontal onset involving the middle and superior frontal gyri; right anterior hippocampal and amygdalar involvement |
| 3 | 25 | M | L | White | 11 | Left dorsomedial prefrontal onset |
| 4 | 54 | M | R | White | 11 | Right temporal onset involving the inferior temporal pole and amygdalohippocampal complex |
| 5 | 41 | M | Ambidextrous | White | 5 | Right temporal onset |
| 6 | 14 | M | L | White | 7 | Left mid-to-posterior cingulate onset |
| 7 | 65 | F | R | White | 10 | Left temporal/hippocampal onset |
| 8 | 42 | F | R | White | 23 | Left precentral and temporal onset involving the lateral temporal pole and anterior hippocampus |
| 9 | 41 | F | R | Black or African American | 8 | Bilateral/multifocal temporal involvement, including the left perirhinal region and right temporal pole |
| 10 | 60 | F | L | White | 14 | Left frontal onset |
| 11 | 13 | M | R | Black or African American | 6 | Left focal motor seizure onset |
| 12 | 18 | F | R | White | 17 | Left mesial temporal lobe focal epilepsy |
| 13 | 16 | F | L | White | 14 | Left parahippocampal and fusiform gyrus onset |
| 14 | 36 | F | R | White | 10 | Left temporal onset involving the posterior hippocampus |
| 15 | 53 | F | L | White | 6 | Left frontal-onset focal epilepsy |
| 16 | 30 | F | R | White | 7 | Poorly localized focal epilepsy; cingulate onset hypothesized |
| 17 | 29 | M | R | White | 7 | Right temporal onset involving the amygdala and anterior and posterior hippocampi |
| 18 | 25 | M | R | Black or African American | 21 | Unknown seizure onset with bilateral temporal epileptiform activity |
| 19 | 27 | F | R | White | 7 | Left hippocampal onset associated with a left temporopolar encephalocele |
| 20 | 33 | F | R | White | 7 | Left anterior hippocampal onset with multifocal temporal epileptiform activity |

N/A: not available. In the Sex column, 'F' stands for 'female' and 'M' stands for 'male'. In the Handedness column, 'L' stands for 'left' and 'R' stands for 'right'. Monitoring duration (days) was calculated from the implantation date through the explantation date, inclusive.

**Movie S1**. Experimental setting for intracranial recording during taVNS.
